# Handwritten Digit classification with neural cultures is influenced by neural architecture, network dynamics, and decoding methods

**DOI:** 10.64898/2026.08.10.743829

**Authors:** Alon Loeffler, Forough Habibollahi, Kwaku Dad Abu-Bonsrah, Azin Azadi, Candice Desouza, Hui Wen Chan, Yusei Nishi, Johnson Zhou, Finn Doensen, Hideaki Yamamoto, Brad Watmuff, Brett J. Kagan

## Abstract

As silicon-based computing approaches fundamental physical limits, neurocomputing offers an energy-efficient alternative by leveraging the intrinsic non-linear dynamics of biological systems. To harness these dynamics, it is vital to understand the structure-function relationship governing how neural cultures process complex spatio-temporal information and how to appropriately decode the resulting neural electrophysiological activity. We investigated this utilizing a closed-loop electrophysiology platform, the CL1, to implement reservoir computing in human iPSC-derived neuronal networks. To systematically evaluate the variables driving neurocomputational capacity, we explored how cellular composition (cortical vs. hippocampal lineages), and the physical architecture (unstructured monolayers, 3D neural organoids, and modular networks confined by microfluidic devices) influenced electrophysiological properties and interacted with different decoding methodologies. Using a spatio-temporal version of a handwritten digit pattern recognition task (MNIST), we analyzed how these biological and analytical factors influenced classification accuracy. To ensure robust interpretation this required us to first demonstrated that reservoir computing decoding methods require strict artifact control and trial-based cross-validation to distinguish network computation from artifactual signal separability or temporal data leakage. Applying this validated frequency-domain pipeline, we suggest a clear functional hierarchy where structural modularity acts as a vital functional regularizer. Modular cortical cultures significantly outperformed unconstrained monolayers and organoids on MNIST. Furthermore, decoding frequency information from raw signals proved superior to typical time-bin decoding implementations. These findings establish that maximizing the computational potential of Synthetic Biological Intelligence, while avoiding false positives, requires a synergistic optimization of cellular identity, structural governance, and rigorous decoding logic. In doing so, this work provides a critical base establishing the criteria under which to evaluate neurocomputing implementations.

## 1 Introduction

A key challenge in Synthetic Biological Intelligence (SBI) and neurocomputing is understanding the drivers of biological neural networks as highly non-linear, multidimensional, dynamical systems. The behavior of these dynamic systems emerges from the interplay between (1) their internal processes and (2) the external structural forces acting upon them [1, 2]. Neural systems present a unique challenge because their external environment encompasses both the informational domain in which they are embodied and the physical constraints dictated by their geometry and topology. While these internal and external factors operate across multiple scales and domains, here we focus on cell type, information decoding, and the macroscale geometry and topology of the neural cultures.

To account for the diverse internal processing capabilities of different cell types, previous works exploring neurocomputing have used predominately cortical cells generated either from primary rodent cultures [3–5] or differentiated from human induced pluripotent stem cells (iPSCs) [6–8]. Other work has explored the use of hippocampal cultures from primary rodent preparations [9–11]. However, only recent advances in differentiation methods have facilitated the development of all major subtypes of hippocampal cells from iPSCs [12]. This means that no previous work has been able to directly compare how human cortical cultures may differ in their response to human hippocampal cultures. Both cortical and hippocampal cells are known to be involved in several critical cognitive processes. The cortex is critical in most higher order cognitive functions, including language, reasoning, inference tasks, and information integration [13–16]. The hippocampus is also a vital region, involved in predictive modeling, adaptability, and memory consolidation, among other functions [17–20]. While it is difficult in vivo to separate out the cell type, it is possible to explore the differences between cortical and hippocampal cells by differentiating iPSCs into both cell types.

To consider how the physical structure of a neural culture impacts information processing, its geometry and topology can be controlled. Modular networks (i.e. networks typically prepared by culturing neurons in microfluidic devices fabricated from polydimethylsiloxane (PDMS)) can be designed to synthetically enforce the geometry and topology of a culture to increase modularity in a biological neural network. Modularity has been shown to increase the separation of dynamical trajectories and improves the performance on reservoir computing (RC) pattern classification tasks [21–23].The intricate activity patterns seen in these networks are usually driven by clustered, modular structures. This organized clustering may allow the network to process information much more effectively than randomly connected networks [24, 25]. Beyond simply increasing dynamic complexity, modularity drives functional specialization by segregating the network into distinct units [26]. By clustering connections locally, modularity has also been shown to facilitate structured synchrony of neural cultures, enhancing their sensitivity to external asynchronous perturbation without triggering global signal saturation [27, 28].

Information is typically inputted through patterned electrical pulses [3, 29]. RC offers a powerful, controlled framework for understanding the fundamental drivers of activity changes. In RC, inputs are fed into a high-dimensional dynamical system called the reservoir [30, 31]. The internal weights of the reservoir are not trained and are assumed to be fixed (a tentative assumption, especially in the case of living neural networks) [32]. Instead, the reservoir’s inherent non-linear dynamics project the input signal into a high-dimensional state space. Crucially, successful RC relies on the principle of ‘fading memory,’ where the network retains a transient, temporal trace of recent inputs within its ongoing reverberations. This intrinsic autocorrelation integrates sequential information, allowing a simple linear classifier trained on the outputs to separate the data [33]. Recently extended to biological neurons [7, 21, 34], RC provides an ideal computational paradigm to evaluate biocomputing capabilities of neuronal systems.

Despite these promising results, modular cultures remain underexplored and there remain questions as to their robustness and relative functional benefits over other culture types. To resolve this uncertainty, this work aims to explore how performance varies between cortical and hippocampal cells across different physical structures, including monolayers, organoids and 60-well modular bioengineered networks. With these preparations, our aim is to implement a spatio-temporal RC task (Figure 1) using the classic MNIST hand-written digit dataset. Once this signal is processed by the neuronal network, its resulting activity must be read out to evaluate performance. Rather than treating the MNIST decoding pipeline as fixed, we apply to it several decoding mechanisms on electrophysiological responses from range of neuronal network types and structures.

**Fig. 1.**
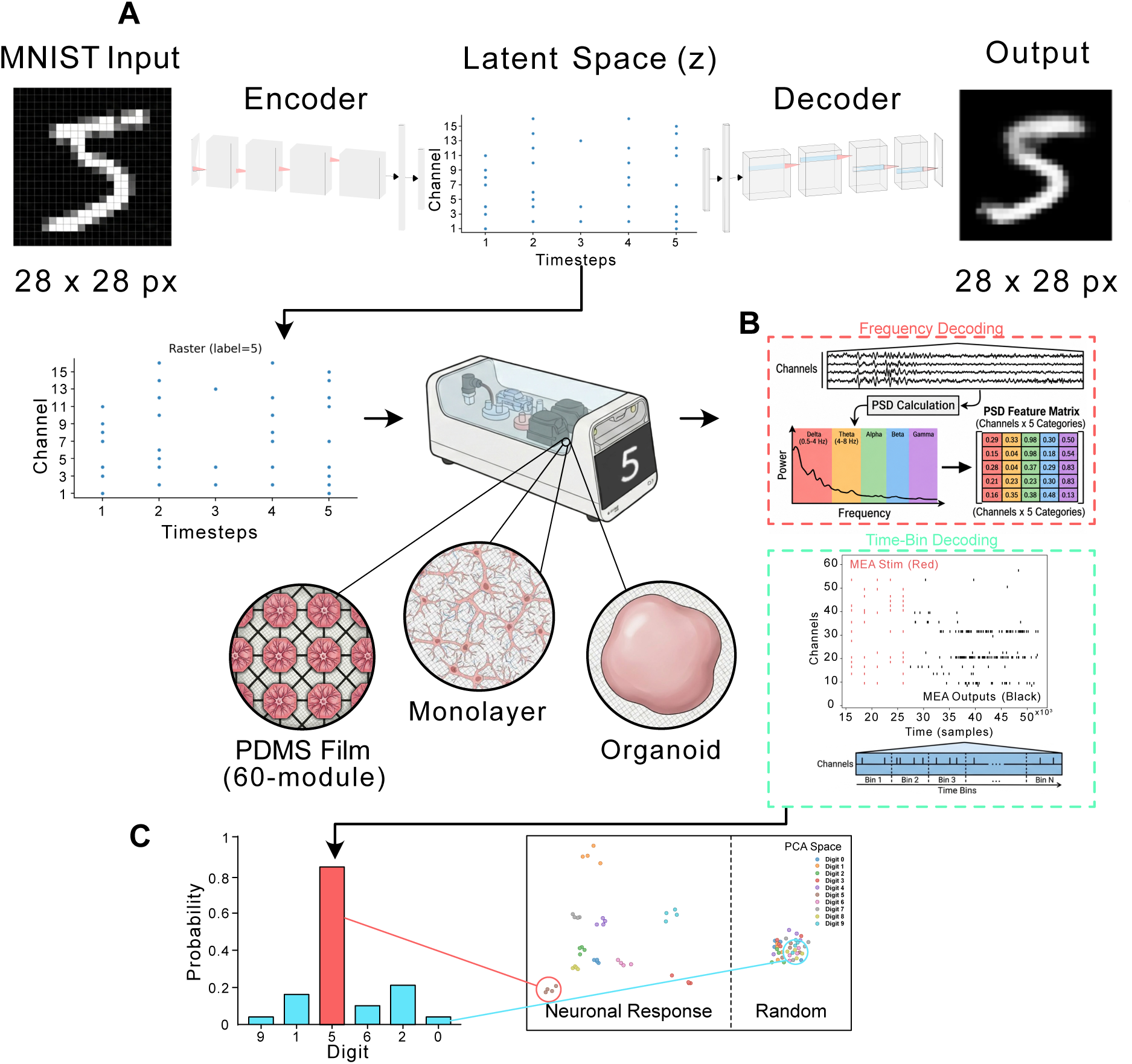
Summary of the RC task and decoding mechanisms compared. **A** Conversion of MNIST digit to spatiotemporal spike-train signal via a spiking autoencoder. An image (28 × 28 pixels) was fed into a multi-layered convolutional neural network (CNN) with leaky integrate-and-fire (LIF) neurons. The latent space of the autoencoder was constructed to represent a spatio-temporal spike-train, validated by reconstruction of the image via a decoder. After training the autoencoder for 100 epochs, a reconstruction loss of ~0.2 was achieved. The latent space of the final epoch was then presented as a sequence of stimulations to a CL1 containing cell cultures on microelectrode arrays (MEAs), in a spatio-temporal MNIST classification task with different neuronal culture setups (PDMS Film 60-module, Monolayer and Organoid). **B** Different decoding mechanisms - Frequency (red) and Time-Bin (cyan) based decoding. **C** Classification of the predicted digit after decoding, and an example PCA space (1st and 2nd PCs) for Neural responses allowing for higher classification accuracy than Random responses.

While RC studies typically report a single accuracy figure per reservoir configuration, comparatively little attention has been paid to how that result depends on the decoding pipeline applied to a given set of recordings. Decoding choices, including which features are extracted from the raw signal (e.g., discretised spike counts versus frequency-domain content), which portion of the response window is analyzed, and how the data are partitioned for cross-validation, are frequently treated as implementation details rather than as variables that can themselves determine whether a reported accuracy reflects genuine neural computation. This is a particular concern for biological RC, where variables such as stimulation artifacts and high-dimensional feature spaces can each independently inflate apparent performance [35–37].

Our investigation is designed to disentangle the drivers of biological information processing by systematically exploring the intersection of biological identity, physical topology, computational decoding, and underlying network dynamics. Specifically, we conduct a moderation analysis utilizing an activity index, a criticality index, and a functional connectivity index to pinpoint the fundamental drivers of performance. We are guided by three core research questions:

**(RQ1)** How does cellular composition (e.g. cortical versus hippocampal) and the physical architecture of the reservoir (e.g. 2D monolayers compared with 3D organoids or bioengineered microfluidics) influence the intrinsic capacity of neural networks to categorize spatio-temporal MNIST patterns?

**(RQ2)** How do the electrophysiological and fundamental network states (i.e., activity, criticality, and functional connectivity) dictate computational performance?

**(RQ3)** How do varying decoding mechanisms (e.g. frequency-domain features vs. temporal spiking vectors) interact with electrophysiological and fundamental network states to modulate the observed performance of the networks?

Rather than hypothesizing that any single variable is dominant, we benchmark these combinations against shared core controls (shuffled output from the neural cultures, a cell-free saline dish, chance, and synthetic random spike trains) to establish which combination of identity, architecture, and decoding pipeline yields accuracy estimates that are both reliably above chance and robust to known confounds. Additional control tests explored spiking neural networks and the capability to cross compare between groups with different activity levels and decoding methods. To enable these experiments, we used the CL1 system, a purpose built MEA platform designed to allow rapid iteration of tightly controlled and reproducible electrophysiological interactions with neural cultures. The CL1 system provides high-level programmability of in vitro neural cells via simple Python APIs [38–40]. This, in turn, enables rapid and iterable development and testing of SBI applications previously unavailable in neurocomputing. Here, we explore the implementation of an open-loop RC approach to leverage the inherent non-linear dynamics of neuronal cultures via the Cortical Labs Application Programming Interface (CL API) [39]. As such, we do not aim to explore learning in these cultures, as no feedback is provided and no change over time is explored. To assess whether our findings generalize across biological substrates, we evaluated each decoding mechanism on recordings from unstructured monolayers, 3D organoids, and microfluidic-modularized (PDMS) networks, spanning both cortical and hippocampal cell identities.

## 2 Results

### 2.1 Generating Neural Cultures

Neural cells derived from human induced pluripotent stem cells (hiPSCs) recapitulate key functional features of in vivo neural systems, including network activity, rudimentary learning capacity, and cellular diversity. Among various brain regions, the hippocampus and prefrontal cortex (PFC) form a core functional axis connected through direct and indirect pathways [41]. The hippocampus, a central hub for cognition and memory, influences decision making, goal directed behaviour, working memory, relational and sequential processing, pattern completion, and contextual integration [17, 42–44]. Differential specialisation along the hippocampal (Hippo) axis supports distinct computational roles, with dorsal circuits contributing primarily to spatial and cognitive functions and ventral circuits associated with emotional and stress related processing [17, 42–44]. The organization of Hippo–PFC connectivity therefore enables bidirectional modulation of information flow and underlies the encoding, retrieval, and storage of information in context dependent domains [41].

To evaluate the generality of the decoding-mechanism comparisons that follow across diverse biological substrates, we generated three types of neural cell cultures on MEAs: strongly interconnected monolayer cultures, highly modular cultures in PDMS films (60-module device), and 3D organoids (Figure 2A). These three groups all generated morphologically robust networks consistent with their respective scaffolds (Figure 2B), while exhibiting characteristic electrophysiological profiles (Figure 2C) and high-fidelity waveforms (Figure 2D) as captured by the CL1 MEA system. Cellular compositions for the monolayer, organoid and 60-module device group included either cortical or hippocampal cells.

**Fig. 2.**
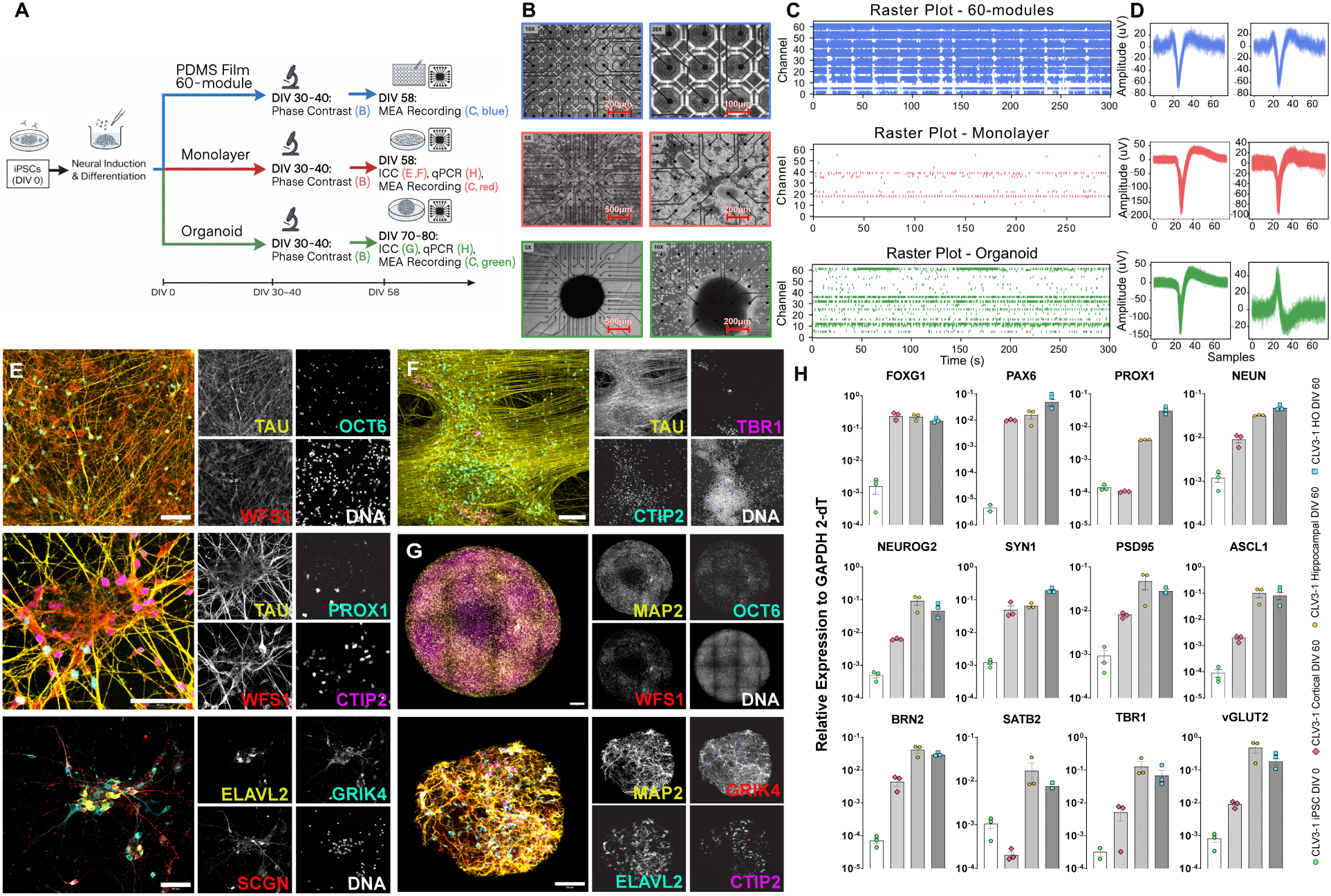
Neuronal cultures. **A**: Schematic illustration of differentiation, plating and recording protocol for each cell structure (60-module device in blue, Monolayer in red, Organoid in Green). Letters in the schematic refer to subplot headings in the current figure. **B**: Microscope images of Top: Modular PDMS Film 60-module device. Each well is connected only to its eight closest neighbors via micro-fluidic channels which encourage axonal growth from well to well. Middle: Monolayers. Bottom: Organoids. **C**: Example 5 minute raster plots for each structure type. **D**: Example waveforms (3 ms cut-out). **E**: Immunostaining analysis after long-term culture shows matured TAU^+^ neurons constituting distinct hippocampal subfield fates: OCT6 and WFS1 (CA1), CTIP2 (CA1 and DG), PROX1 (DG), ELAVL2, GRIK4 and SCGN (CA3). Scale bar is 50 µm. **F**: Immunostaining analysis after long-term culture shows matured TAU^+^ neurons constituting CTIP2^+^ (cortical layer V) and TBR1^+^ (cortical layer VI). Scale bar is 100 µm. **G**: Immunostaining analysis of Hippo Organoids shows matured MAP2^+^ neurons constituting distinct hippocampal subfield fates: OCT6 and WFS1 (CA1). Scale bar is 50 µm. **H**: qRT-PCR confirms the expression of proneural markers and markers of cortical and hippocampal neurons at DIV 60 (n = 3 independent experiments each, data are mean ± SEM).

To reconstruct aspects of these neural systems in vitro, we confirmed the specification of major hippocampal subfield neurons in differentiated cultures by immunostaining. In cultures which expressed the mature neuronal marker TAU, we observed CA1 neurons expressing WFS1, OCT6, and CTIP2, dentate gyrus (DG) neurons expressing PROX1 and CTIP2; and CA3 neurons marked by GRIK4, ELAVL2, and SCGN (Figure 2E, top, middle, and bottom respectively). We further validated the identity of differentiated cortical neurons, and found cells expressed cortical layer V (CTIP2^+^) and VI (TBR1^+^) markers (Figure 2F). In Hippo Organoids at DIV100, immunostaining demonstrated the presence of GRIK4^+^/ELAVL2^+^ CA3 neurons and WFS1^+^ CA1 neurons (Figure 2G). Transcriptional profiling at DIV0 and DIV60 corroborated these observations, showing expression of proneural genes and markers characteristic of both cortical and hippocampal neuronal lineages (Figure 2H).

Together, these findings establish a framework for generating and characterising hiPSC derived neural systems capable of modeling region specific identities and network level properties.

### 2.2 Electrophysiology and Criticality

Figure 3 summarizes the overall electrophysiological activity metrics recorded for each cell culture configuration. Eight key metrics were analyzed: Mean Firing Rate (Hz; A), Variance of Firing Rate (Hz^2^; B), Max Firing Rate (Hz; C), Mean Inter-Spike-Interval (ISI) (s; D), Variance of ISI (s^2^; E), Deviation from Criticality Coefficient (DCC) (F), Branching Ratio (G) and Shape Collapse Error (H). Full statistical comparisons across all cell types are detailed in the Supplementary Materials.

**Fig. 3.**
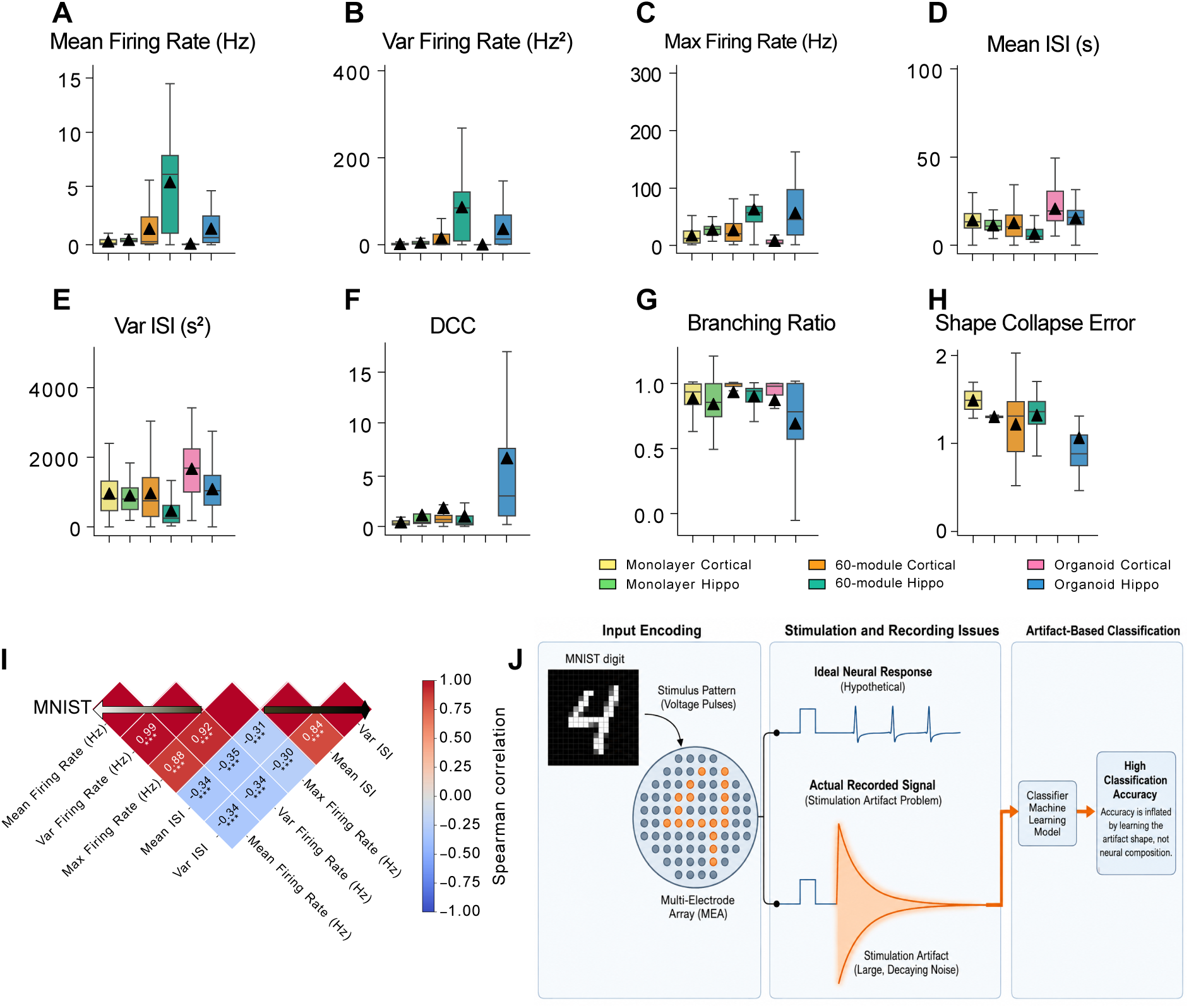
Electrophysiological and criticality metrics across different cell culture architectures. Spontaneous culture-level activity is compared across diverse cellular configurations using boxplots. Culture ages ranged from DIV 58 for Monolayers to up to DIV 80 for Organoids. A total of 53 cultures were recorded both baseline (spontaneous) and playing MNIST (see Supp Table S1 for full details). **(A–C)** Spiking statistics, including Mean Firing Rate, Variance of Firing Rate, and Maximum Firing Rate. **(D,E)** ISI metrics, showing Mean and Variance of ISI. **(F–H)** Network criticality metrics, specifically the DCC, Branching Ratio and Shape Collapse Error. For all plots, boxes represent the interquartile range (IQR) and median, whiskers extend to 1.5×IQR, and black triangles denote the mean. The metrics highlight distinct spontaneous activity profiles, demonstrating elevated activity and near-critical dynamics in modular networks compared to unstructured monolayers and organoids. **I** Heatmap comparing correlations in electrophysiological activity during the MNIST task. **J** Neuronal activity can be masked by stimulation artifacts if not properly blanked. This affects both functional connectivity of the cultures as well as classification accuracy.

Overall task-related network activity was most pronounced in the structured modular networks, particularly the 60-module Hippo cultures. As shown in Figure 3A–C, the 60-module Hippo group showed the highest median values and greatest upper ranges across all spiking metrics (Mean, Variance, and Maximum Firing Rates), indicating a state of heightened overall activity, with large fluctuations. The 60-module Cortical and Organoid Hippo cultures also exhibited moderate activity levels. In contrast, many trials implementing unstructured monolayer configurations (Cortical or Hippo) displayed consistently low firing rates, with Organoid Cortical cultures showing the lowest overall spiking activity. We acknowledge that organoids have a different architecture as seen in 3B where active areas of the culture may not reach every single electrode, but as can be seen in 3C, organoids typically had more active channels that monolayers despite this feature allowing a useful comparison of the activity across the network relative to the architecture. See 5.2.1 for our selection criteria for cultures and recordings.

Interestingly, the 60-module Hippo group exhibited the lowest mean and variance for its ISI (Figure 3D, E). This suggests that its activity is characterized by highly regular firing patterns, while other groups such as the Organoid Cortical, may be more characterised by irregular or “bursty” firing patterns. In terms of the criticality metrics, the Organoid Hippo group diverged from the rest (Figure 3F–H): although it had the lowest Shape Collapse Error, it demonstrates a markedly higher DCC, a noticeably lower Branching Ratio (dropping to a mean near 0.7 compared to the near-critical 1.0 observed in the other groups). These combined results suggest that this group operates in a distinct, likely subcritical, dynamical regime compared to the other evaluated conditions, despite having similar temporal avalanche profiles. Contrastingly, all other groups showed evidence of criticality, although with high variability, with median points clustered around 0 for DCC, 1 for Branching Ratio, and *>* 1 for Shape Collapse Error. We next wanted to test multicollinearity between electrophysiological features during a task, rather than just at baseline spontaneous activity. To do so, we compared electrophysiological activity metrics recorded during the MNIST task. Figure 3I shows strong correlations between metrics, suggesting multicollinearity between those features.

### 2.3 Functional Connectivity

To characterize global organization and the degree of network integration, functional connectivity was estimated from multi-channel spike recordings by first coarse-graining spike times into discrete temporal bins of 1 s duration. For each channel, spike counts were computed within each bin to obtain a time-resolved representation of population activity. Pairwise functional connectivity was then quantified using the Pearson correlation coefficient between binned spike-count time series across all channel pairs, yielding a weighted connectivity matrix. To focus on robust interactions, connections were thresholded by absolute correlation magnitude (|*r*| ≥ 0.4). Self-connections were excluded. The resulting thresholded matrix was interpreted as an undirected weighted graph representing functional interactions among channels.

From this graph, we extracted several summary network measures that provide a compact description of the strength, segregation, and integrative properties of the functional networks. 60-module Hippo cultures exhibited the highest Total Edge Weights and Average Edge Weights (Figure 4A, B). In terms of local and mesoscale organization, the 60-module Hippo networks demonstrated the highest average weighted clustering coefficient, reflecting strong local segregation [45] (Figure 4C). Conversely, Modularity (derived from Louvain community detection) was highest in both the Organoid Hippo and 60-module Hippo networks, highlighting their strong mesoscale community structure (Figure 4D). Finally, both the 60-module and Organoid Hippo networks displayed high maximum weighted betweenness centrality, indicating prominent network hubs and critical information flow bottlenecks (Figure 4E). These spatial topologies and hub locations are visualized across the different culture architectures in Figure 4F–L, mapping node degree and betweenness centrality to physical electrode locations. Together, these topological metrics indicate that both physical microfluidic confinement and hippocampal cell identity promote highly organized, modular networks centered around strong local hubs.

**Fig. 4.**
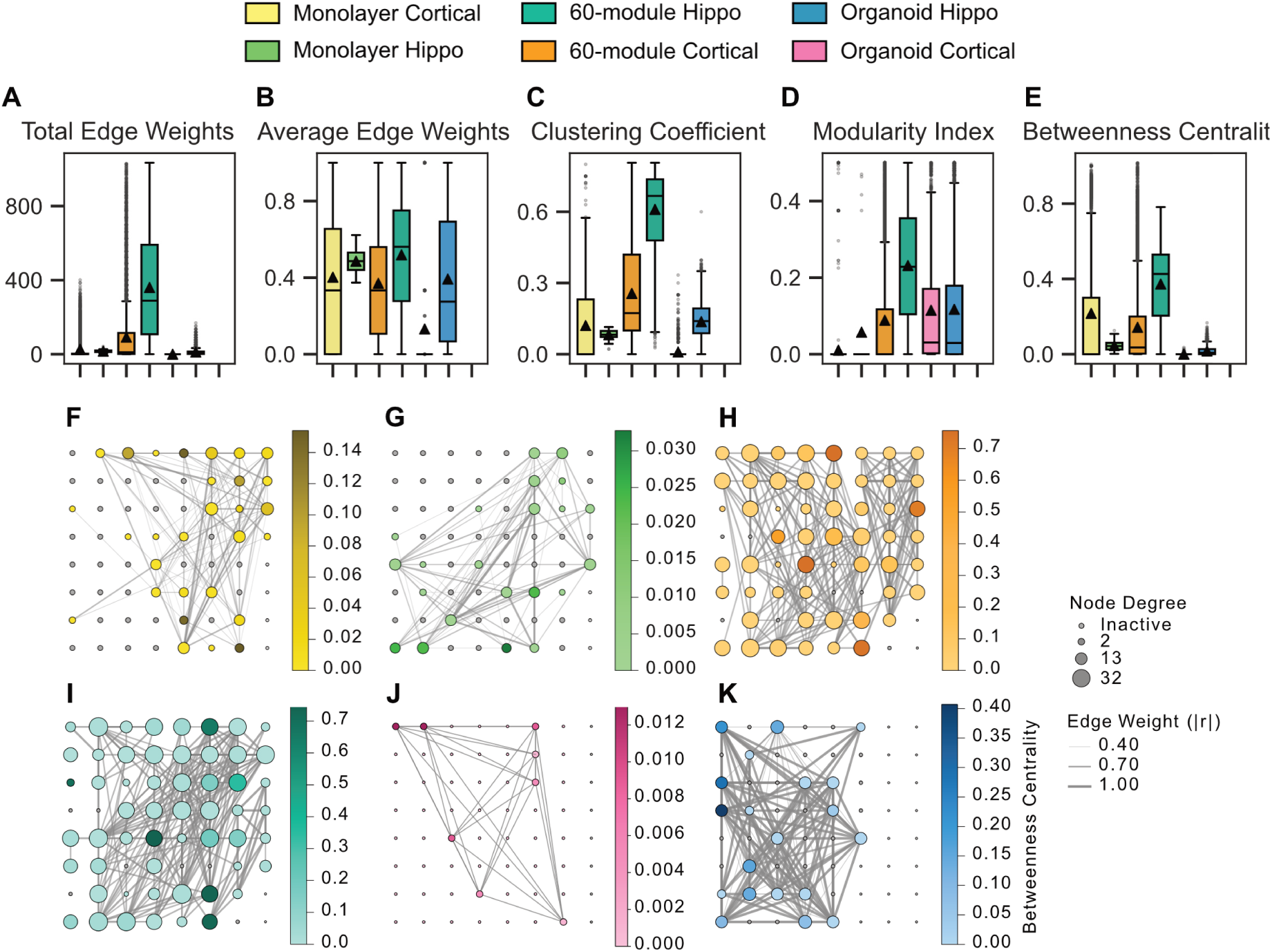
Functional connectivity and network topology across culture architectures. Culture-level network graph metrics are compared across diverse cell configurations using boxplots with overlaid individual data points **(A–E)**. Functional connectivity was measured as thresholded Pearson correlations (|*r*| ≥ 0.4) between electrodes. Boxes show the interquartile range and median, whiskers indicate 1.5×IQR, and black triangles denote the mean. **(F–K)** Representative spatial functional connectivity maps for each culture category. Nodes represent active electrodes, with size indicating node degree and color mapping to betweenness centrality. Edge thickness reflects connection weight, illustrating the distinct topological integration and hub formations specific to each architecture.

### 2.4 Stimulation Artifacts and Classification Errors

Precise bidirectional interrogation of neural circuits is frequently confounded by stimulation artifacts; transient, high-amplitude voltage excursions that obscure neuronal activity immediately following the stimulus [35]. Arising from the capacitive discharge at the electrode-electrolyte interface, these artifacts often saturate recording amplifiers, creating a temporal ‘blind spot’ (typically 2–10 ms) that masks critical short-latency synaptic responses [36]. While software-based subtraction and hardware blanking offer partial mitigation, residual non-linearities have historically limited the ability to detect direct network recruitment with high fidelity [37]. Consequently, these artifacts must be blanked prior to analysis to prevent high-amplitude transients from confounding classification accuracy. Failure to mitigate these artifacts not only obscures the underlying physiological signal, rendering the intrinsic neural dynamics undetectable (see Figure 3J and Supp Figure S5), but also can lead to false-positive activity detections that represent a systematic error that can confound experiments. This may result in inflated electrophysiology activity, functional connectivity and most worryingly, classification performance driven by the geometric distinctness of the stimulation artifacts rather than the non-linear computational capacity of the neuronal networks. To ameliorate this issue, we employed both a hardware and software solution to blank stimulation artifacts for 2 ms post-stim, ensuring no residual input signal in read-out measures. We also include results which include partial stimulation artifacts to elucidate this concern. Furthermore, as explained in Methods section 5.3, any stimulation artifact waveforms that may have been missed by blanking were removed by spike sorting and clustering.

However, blanking stimulation artifacts alone is not sufficient to confirm that any decoded response reflected actual changes in neural culture activity patterns. We therefore implemented additional controls, as described below.

#### 2.4.1 Importance of proper controls

During the Time-Bin classification task (see Section 2.5 for details), we evaluated a common readout in RC, logistic regression with L_2_ regularization (Ridge) for classifying reservoir states. Our control experiments using randomly shuffled data revealed a critical vulnerability of high-dimensional feature spaces: when there is high network activity, the number of bins used to partition the data drives overfitting more strongly than the readout algorithm itself. Specifically, sorting data into a high number of bins forced the feature-to-sample ratio so high that it yielded near-perfect, spurious accuracy (up to 99%) by exploiting random correlations (spurious separability; see Supp Figure S6). Importantly, such accuracy was seen mainly in highly active and highly connected networks (see Figure 5K&L), meaning the implementation of appropriate controls to identify and mitigate these issues even more important. We detail our methods in Section 5.4.1. To combat such overfitting issues, we utilized L_2_-regularized logistic regression alongside a valid binning strategy. The L_2_-penalty (Ridge regularization) introduces a cost term proportional to the square of the magnitude of the model’s coefficients. This regularization effectively penalized and shrank the weights for non-informative, noisy, or highly correlated features, preventing the classifier from overfitting to the training data and ensuring the model relied only on the most robust predictors [46].

**Fig. 5.**
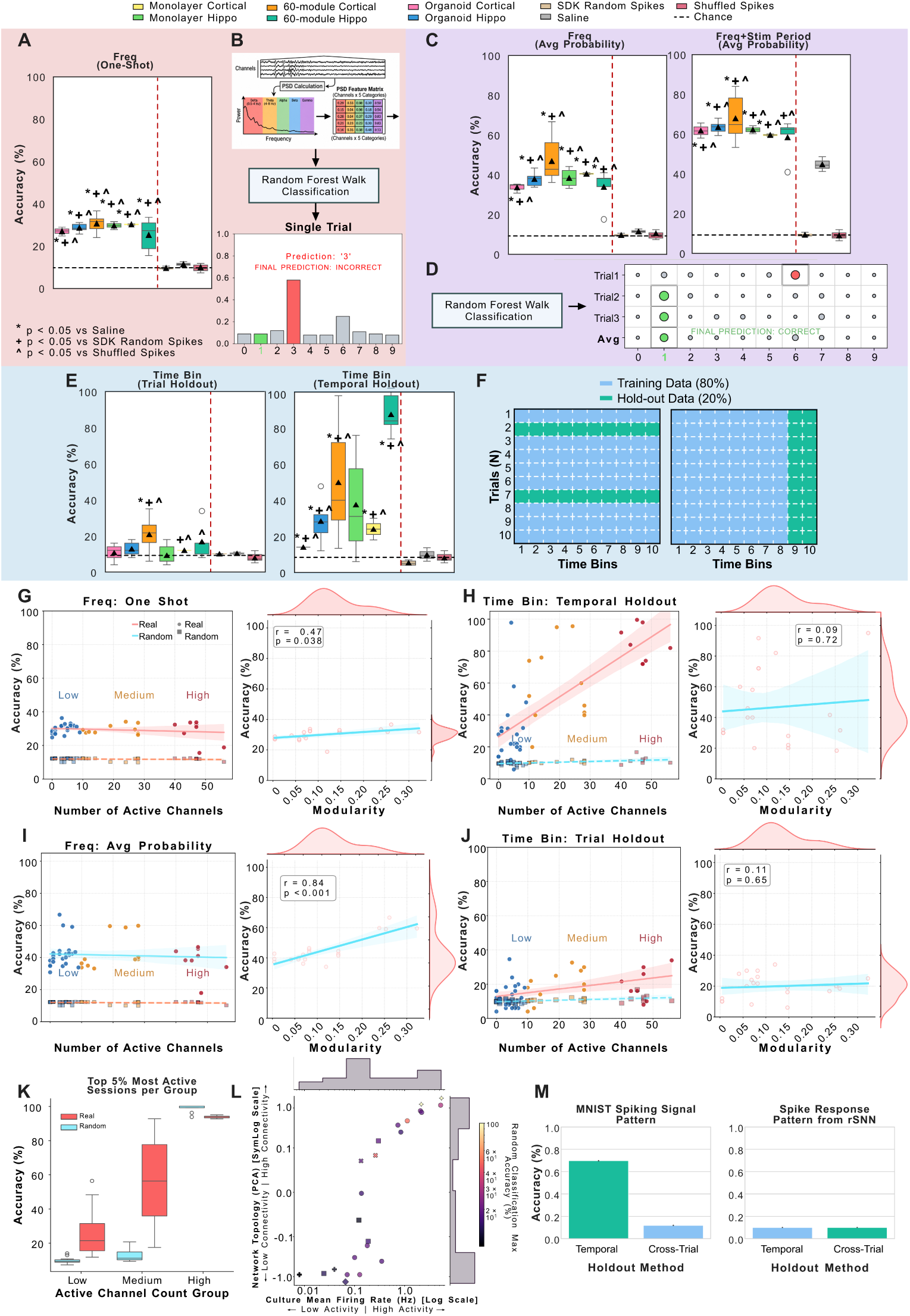
Classification Accuracy Explored by Cell Type, Decoding Methodology, and Functional Connectivity. **(A–B, Red)** Frequency-domain one-shot classification per digit trial. **A:** Accuracy by cell type and controls: Monolayer Cortical (*n* = 2), Monolayer Hippo (*n* = 8), 60-module Cortical (*n* = 22), 60-module Hippo (*n* = 5), Organoid Cortical (*n* = 3), Organoid Hippo (*n* = 5). Dashed lines: horizontal = chance (10%); vertical red = separates neurons (left) from controls (right). **B:** Calculation schematic. Significance (t-test vs. controls): ∗*p <* 0.05 vs. Saline; +*p <* 0.05 vs. SDK Random Spikes; ^*p <* 0.05 vs. Shuffled Spikes. **(C–D, Purple)** Multi-trial average probability frequency-domain decoding. **C:** Results without (left) vs. with (right) the stimulation window. Including the stimulation window drastically alters apparent accuracy and variability; elevated saline control accuracy under “with Stim” indicates partial contamination by stimulation artifacts. **D:** Multi-trial calculation schematic. **(E–F, Blue)** Time-bin holdout decoding. **E:** Trial holdout (left) vs. time holdout (right). **F:** Train-test split schematics for trial holdout (80% train / 20% test trials; left) and time holdout (80% train / 20% test time bins; right). **(G–J)** Network activity and modularity comparisons. Activity is operationalized by active channel count (≥ 75% spike ratio; *>* 1400 spikes): Low (*<* 10), Medium (10–30), High (*>* 30). Left panels compare active channels against frequency decoding (G, I) and time-bin decoding (H, J). Right panels show classification accuracy vs. Modularity metric. **K:** Neuronal vs. randomly shuffled classification accuracy across top 5% low, medium, and high activity groups. **L:** Network topology (y-axis), culture mean firing rate (x-axis), and random classification accuracy (hue). Higher connectivity and activity produce higher random accuracy due to overfitting and bin misalignment. **M:** Cross-trial (*N* -axis) vs. temporal (*T* -axis) holdout classification accuracy. **Left:** The stimulation template control (raw MNIST spatio-temporal spike train) drops from 69.7% to 11.9% in Cross-Trial holdout. **Right:** The rSNN baseline (two RSynaptic neuron layers) maintains stable accuracy across holdouts, demonstrating that biological culture temporal discriminability is unlikely a data-leakage artifact.

To validate that the model captured a true input-output relationship rather than statistical artifacts, we performed permutation tests by randomly shuffling the target labels.The L_2_ regularization prevented the algorithm from simply learning random feature distributions and correctly returned chance-level performance on shuffled data (where the true mapping between inputs and target labels was randomly broken). Ultimately, the convergence of structural feature penalization and permutation testing provides a much more robust and trustworthy measure of the system’s generalized computational capacity. We note that even properly binned, regularized models can produce artifactual results if confounding variables, such as stimulation artifacts or high network activity, are not properly addressed. To mitigate this risk, we incorporated two additional comparators alongside the randomized post-stimulation group: a media-only condition (MEA with no cells) to rule out hardware artifacts, and an in silico benchmark (randomized activity readout) to exclude systematic software artifacts in encoding or decoding.

### 2.5 Classification

#### 2.5.1 Comparing Decoding Mechanisms for MNIST Classification

We used the classic MNIST handwritten digit recognition dataset as a single, well-characterized spatiotemporal classification task on which to benchmark different decoding mechanisms. Digits were converted into spatio-temporal spike trains and delivered to neural cultures via the CL1 system (see Methods). From the exact same set of recordings, we then decoded digit identity using four distinct pipelines (Figure 5A,B and C), benchmarked against shuffled, no-cell (saline), and synthetic random-spike controls:

- **Frequency-domain**: Five-band power spectral density (PSD). Features of the raw signal in a 400 ms window beginning 10 ms *after* the final stimulation pulse, decoded via a Random Forest, with cross-validation performed across trials.
- **Frequency-domain with Stim**: identical to the above, except the analysis window began at stimulation onset, so the response includes the stimulation period itself.
- **Time Bin (Trial Holdout)**: spike-count time-binning approach (66.7 ms bins), decoded via logistic regression, cross-validated across trials.
- **Time Bin (Time Holdout)**: identical features and classifier to Time Bin (Trial Holdout), but cross-validated by holding out time bins rather than trials. This acts to explore not whether the cultures can classify digits, but offer responses that are predictive of missing information.

Full methodological details for each pipeline are provided in Section 5.4. Classification performance for all four mechanisms, plus the shuffled, saline, and synthetic random-spike controls, is summarized in Figure 5A–F.

Wilcoxon signed-rank tests (paired by recording) confirmed that every decoding mechanism produced accuracy significantly above its corresponding shuffled control (all *p <* 0.01). However, the margin above chance varied enormously by an order of magnitude across mechanisms. Time Bin (Trial Holdout), was only marginally above chance (mean 14.0% versus 12.2% for shuffled controls; difference 1.7 percentage points, *p* = 0.0074), and above the 10% chance level for this ten-class task by a similarly small margin. In contrast, Frequency-domain achieved an average accuracy of 39.3%, an improvement of 27.0 percentage points over shuffled controls (*p <* 0.001), and was 23.6 percentage points higher than Time Bin (Trial Holdout) applied to the same recordings (*p <* 0.001).

Holding this feature representation fixed and varying only the cross-validation strategy revealed a substantial effect of holdout choice. Time Bin (Time Holdout) more than doubled the average accuracy of the trial-based holdout applied to identical spike-count features (mean 30.0% versus 14.0%; difference 18.0 percentage points, *p <* 0.001), and increased variability dramatically: the coefficient of variation of accuracy across recordings rose from 0.52 (trial holdout) to 0.74 (time holdout), with individual recordings reaching up to 98% accuracy under the time-based holdout despite a maximum of only 35% under the trial-based holdout on the same data.

Frequency-domain with Stim produced the highest average accuracy of any mechanism (62.5%; 22.7 percentage points higher than Frequency-domain on the same recordings, *p <* 0.001). However, this mechanism was also the only one for which the saline (no-cell) control showed markedly elevated average accuracy: 44.4%, far above both the shuffled baseline (∼10%) and the saline control’s own accuracy under the Frequency-domain mechanism (11.3%). Because the saline dish contains no neurons, this indicates that a substantial portion of the accuracy gain from including the stimulation window reflects decoding of the deterministic stimulation artifact itself, rather than genuine neural computation. The synthetic random-spike control, which is not exposed to real stimulation events, remained near chance under both windowing schemes (9.3% and 8.8%, respectively), confirming that this inflation is specific to stimulus-locked artifact rather than a general property of the decoding pipeline.

Spearman correlations between pairs of decoding mechanisms, computed across individual recordings, ranged from *ρ* = 0.31 to *ρ* = 0.64 (all *p <* 0.03), indicating that while mechanisms broadly agree on which recordings perform better than others, the specific ranking of cultures is not fully preserved when the decoding mechanism changes. 60-module Cortical cultures achieved the highest, or joint-highest, median accuracy under every mechanism tested (e.g., 42.7% under Frequency-domain, *n* = 22), suggesting this finding is directionally robust to decoding choice.

Interestingly, while network activity (measured as number of active channels, *<* 10 channels = low, 10-30 channels = medium, *>* 30 channels = high) showed no difference across Frequency-domain accuracy (Figure 5G–J), higher network activity was positively correlated with higher accuracy for the Time-Bin decoding method.

#### 2.5.2 Comparing Cell Types

When comparing cortical cultures (unlabeled) to hippocampal cultures (labeled “Hippo”) across matching physical architectures, a distinct difference in temporal dynamics emerged. Under the more rigorous, artifact-free Frequency-domain mechanism, cortical cultures generally achieved higher classification accuracies than their hippocampal counterparts (e.g., 60-module Cortical mean 47%, max 66.7%, n = 22 vs. 60-module Hippo mean 34%, max 41.0%, n = 5, see Figure 5A–D). However, across all three architectures (Monolayer, Organoid, and 60-module), the time-based holdout show significantly higher hippocampal accuracy compared to cortical variants. This peaked at a mean of 74% (max 84.0%) for 60-module Hippo, compared to a mean of 51% (ma× 100.0%) for 60-module Cortical (see Figure 5E). This suggests that hippocampal networks may exhibit longer temporal autocorrelations or sustained network reverberations that significantly increase accuracy across temporally adjacent bins when trial boundaries are ignored.

#### 2.5.3 Comparing Cell Structure and Function

Evaluating the impact of physical cell culture architecture on decoding performance revealed a consistent functional advantage for the structurally constrained environments. This can mostly clearly be seen on the artifact-free Frequency-domain pipeline where the 60-module Cortical cultures achieved the highest mean classification accuracy (mean 47%, max 66.7%). Standard Cortical Monolayers followed (mean 41%, max 41.0%, *n* = 2), while Cortical Organoids exhibited the lowest decoding performance among the live biological cultures (mean 34%, max 35.3%, n = 3). A similar hierarchical trend is visible across the hippocampal variants. This is consistent with findings shown in Figure 5G–J, and indicates that higher modularity is correlated with higher classification accuracy for frequency decoding methods. No such relationship can be seen in Time Bin decoding methods. Other functional connectivity metric comparisons can be seen in Supp. Figure S4. Comparisons with various controls are represented by *p*-value markers in Figure 5A,C and E.

#### 2.5.4 Validation of Temporal Holdout

Given spatio-temporal feature tensors with dimensions *N* × *C* × *T* , we distinguish between cross-trial holdout (partitioning the *N* -axis) and temporal holdout (partitioning the *T* -axis). As previously noted in Section 2.5.1, aggregating network responses across multiple trials significantly enhances discriminability by mitigating inter-trial variability. We therefore extend this concept by applying temporal holdout across the *T* -dimension to determine the maximum representative signal obtainable from a single stimulus event.

Because temporal partitioning of time-series data is prone to confounding via data leakage, we implement two in-silico controls to quantify the impact of this strategy on predictive performance: (1) The stimulation template control, which utilizes the raw MNIST spatio-temporal spike-train defined in Section 5.4, and (2) the rSNN baseline, which involves simulating a recurrent spiking neural network (rSNN) response to the same input sequence.

For both controls, predictions are performed using an *L*_2_-regularized logistic regression classifier consistent with Section 2.4.1 and repeated with 200 random seeds. In the stimulation template control, we directly evaluate the separability of the fixed input tensor. For the rSNN baseline, we implement a recurrent architecture composed of two layers of RSynaptic neurons via the snnTorch package [47], where the network processes the stimulation pattern via a series of linear transformations (fc in → rsynaptic1 → fc out → rsynaptic2) to produce a windowed spike response from the MNIST spatio-temporal spike-trains in (1). We evaluate the model’s ability to classify digits based on this simulated activity, using a response window identical to our biological experiments.

We observe (see Fig 5M) that temporal holdout causes a significant increase in predictability for the stimulation template (from 11.9% to 69.7%), yet produces no change in accuracy for the rSNN baseline. This divergence suggests that the enhanced discriminability observed in our biological cultures is not an artifact of the methodology but arises from the inherent spatio-temporal continuity present in the biological neural response. While temporal autocorrelation exists within the bins, our rSNN baseline demonstrates that this autocorrelation does not lead to an artificial inflation of accuracy, thereby isolating the biological ‘feature reinforcement’ as a true property of the neural substrate.

#### 2.5.5 Biophysical Determinants of Reservoir Performance

##### Moderation Analysis of Network State

To determine whether the underlying baseline state of the biological networks influenced decoding performance, we conducted a moderation analysis across seven network activity and complexity metrics (total observations *n* = 360). After standardizing the predictors and applying False Discovery Rate (FDR) correction for multiple comparisons, only the Deviation from Criticality Coefficient (DCC) emerged as a significant moderator of classification accuracy (*F* = 17.54, *p_F DR_ <* 0.001). In contrast, baseline activity metrics (Mean Firing Rate, Maximum Spike Count, and Inter-Spike Interval (ISI) statistics) did not significantly moderate performance (all *p_F DR_* ≥ 0.49).

To confirm that the lack of moderation by the activity metrics was not an artifact of multicollinearity, we performed a Principal Component Analysis (PCA) on the four activity-related features (Mean Firing Rate, Max Spike Count, Mean ISI, and Variance ISI). The first two principal components captured 91.54% of the total variance in network activity (PC1: 66.98%; PC2: 24.57%). However, unified moderation models utilizing these components confirmed the null result: neither PC1 (*p* = 0.427), PC2 (*p* = 0.947), nor a combined interaction model (*p* = 0.315) yielded significant moderation effects.

A parallel PCA was conducted on the criticality index (DCC, Branching Ratio, and Mean Information Entropy), capturing 80.01% of the variance across two components. While PC1 loaded heavily on both DCC and Entropy, neither the unified PC1 (*p* = 0.324) nor PC2 (*p* = 0.970) interaction models were significant moderators. Taken together, these results suggest that the efficacy of the decoding mechanisms is robust against broad fluctuations in general network firing rates, but remains highly sensitive to the specific dynamical regime captured by the DCC metric alone.

A third PCA revealed that overarching network topology is heavily governed by two independent dimensions of constraint. To generate the Topology index, we performed a parallel PCA on the functional connectivity metrics. The first two components captured 83.60% of the topological variance; PC1 (48.86%) was driven primarily by network integration and routing efficiency (clustering coefficient: 0.67; maximum betweenness: 0.64), while PC2 (34.74%) was heavily associated with modularity (0.75) and inversely related to average edge weights (-0.62). The primary axis (PC1) represents *Centralization*, and measures a network’s reliance on strict structural super-hubs and choke points. The secondary axis (PC2) represents *Segregation*, and measures a network’s tendency to fracture into isolated functional communities. By mapping cultures across this 2D state-space, we were able to test exactly how different structural strategies (such as physical hubs versus functional modules) constrain emergent critical dynamics.

##### Mapping a 3D Computational Landscape

To elucidate how macro-scale network properties interact to support biocomputation, we constructed three composite indices - Activity, Criticality, and Topology. The Activity, Criticality and Topology composites were all derived directly from the primary principal components of their respective feature spaces. Figure 6A shows the PCA mappings of all three composite indices.

**Fig. 6.**
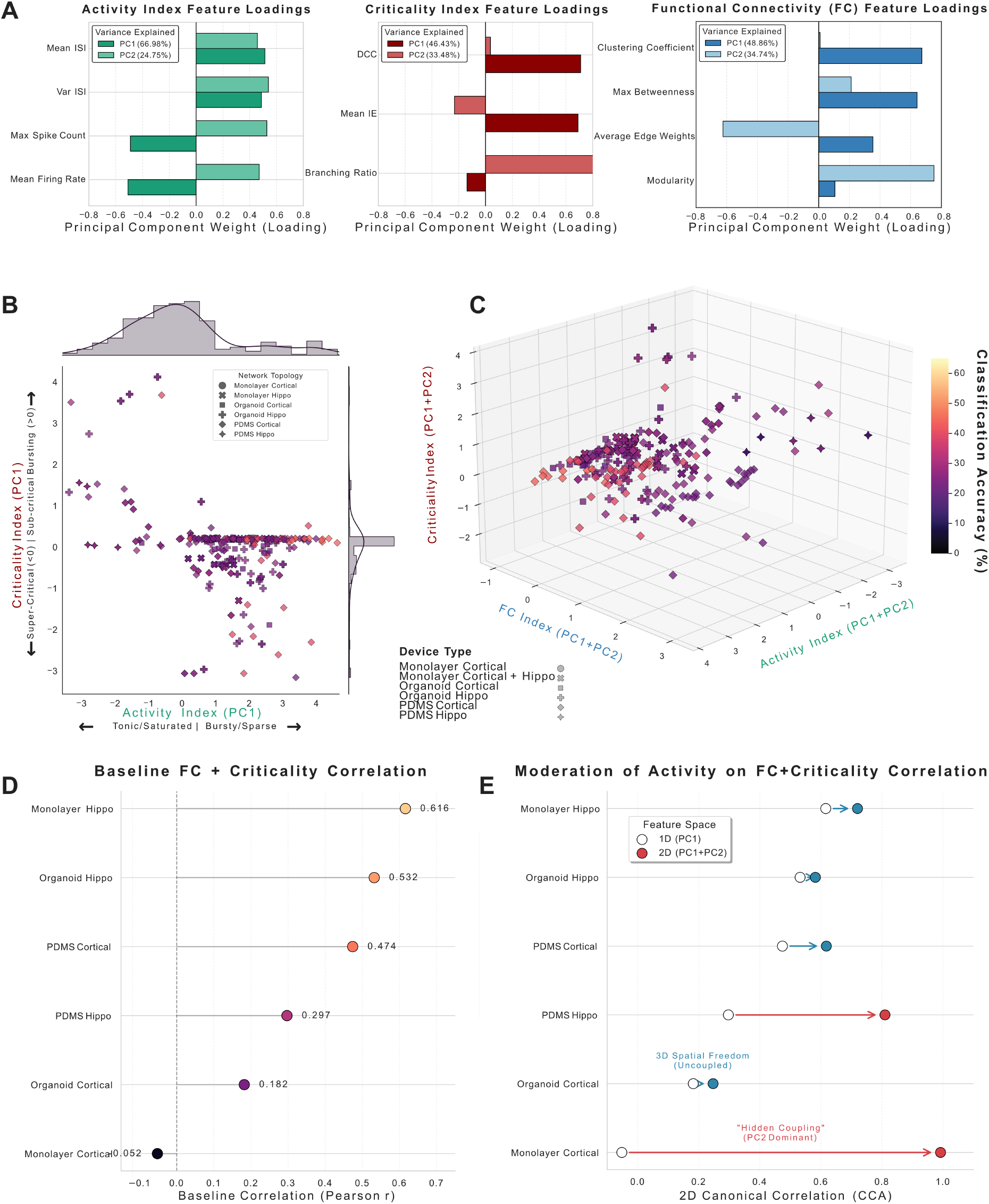
Multi-dimensional state-space mapping and structure-function coupling of in vitro neural architectures. **A** Principal Component Analysis (PCA) feature loadings and explained variance (PC1 and PC2) for the Activity, Criticality, and Functional Connectivity (FC) indices. **B** 2D state-space of network Activity (PC1) versus Criticality (PC1) categorized by device type, featuring marginal distributions along both axes. **C** 3D composite landscape combining PC1 and PC2 for Activity, FC, and Criticality, with data points colored by Classification Accuracy (%). **D** Baseline 1D structure-function coupling (Pearson *r* ) across different device types. **E** Moderation of structure-function coupling, showing the Canonical Correlation Analysis (CCA) shift from a 1D (PC1) to a 2D (PC1+PC2) feature space, highlighting “Hidden Coupling” in 2D monolayers and spatial decoupling in 3D organoids.

To ensure these composite indices accurately reflected distinct biological phenomena, we mapped the principal component (PC) feature loadings back to their underlying physical mechanisms. The resulting state-space coordinates were interpreted as follows:

- **Activity Index:** This index captures the temporal distribution of spontaneous network spiking. PC1 in this feature space weighted Mean ISI (0.51) and Variance ISI (0.49) in direct opposition to the Mean Network Firing Rate (-0.51) and Maximum Spike Count (-0.49). Consequently, this axis functions as a continuum between tonic, high-throughput firing and bursty, irregular activity characterized by long, highly variable inter-spike intervals. By aggregating this with PC2, the resulting composite serves as a unified proxy for the network’s overall temporal structure, with higher scores reflecting networks capable of producing distinct, high-contrast bursts of activity rather than undifferentiated, tonic background firing.
- **Criticality Index:** This index quantifies the dynamical distance from a critical state. PC1 was driven by the DCC (0.71) and Mean Information Entropy (0.69), while PC2 was dominated by the Branching Ratio (0.97). As the DCC measures the deviation from criticality, a composite score near 0 indicates a network operating at the optimal tipping point between order and chaos. Conversely, increasing absolute scores denote a progressive departure from this state, signaling that the network has been pushed into either sub-critical (signal extinction) or super-critical (runaway excitation) dynamical regimes.
- **Topology Index:** This index quantifies the structural integration and routing efficiency of the functional connectivity graph. PC1 loaded primarily on the Clustering Coefficient (0.67) and Maximum Betweenness Centrality (0.64), characterizing networks with dense local interconnectivity and prominent, highly traversed routing hubs. PC2’s heavy weighting of the Modularity Index (0.75) indicates that a maximized Topology composite describes a highly organized, “small-world” architecture, where modular processing hubs are integrated by efficient global pathways, distinguishing these networks from randomly connected syncytia.

We initially compared the impact of Criticality and Activity indices (Figure 6B) on classification accuracy (specifically, Frequency-domain classification). This revealed a large proportion of cultures in the critical (around 0) regime and bursty/sparse activity performed better. Next, we added the Topology index. By mapping each culture into the 3D state space defined by these indices (Figure 6C), distinct clusters emerged that correlate with both physical architecture and decoding efficacy. The best performing networks, 60-module Cortical, occupied a unique region characterized by maximized topological integration (0.919) and a high Activity Index (1.376). We interpreted this as an indication that PDMS-based constraints successfully cultivate networks that trade high-frequency, tonic background noise for precise, high-contrast temporal bursts or specified activation, priming them for stimulus-locked computation.

Conversely, unstructured Cortical Monolayers occupied a functionally impoverished region. While their clustering near 0 on the Criticality axis (0.053) suggests they operate in a critical regime, they suffer from severely reduced topological complexity (0.116). This indicates that while near-critical dynamics are a necessary condition for biocomputation, they are insufficient in the absence of the high-contrast activity patterns and structural scaffolding provided by modular architectures.

Finally, hippocampal variants demonstrated a divergent state configuration. Hippo Organoids, for example, exhibited high activity (0.957) and moderate topological integration (0.462), but displayed the most significant departure from the critical regime (Criticality: 1.235). This hyper-deviated, structurally loose state aligns with our earlier decoding results. The super-critical dynamics characteristic of these hippocampal networks likely foster the sustained, stimulus-independent reverberations and subsequent fading memory dynamics that account for their higher decoding accuracies in time-binned analyses.

##### Multi-Dimensional Structure-Function Coupling

We initially evaluated structure-function coupling strictly along the primary Centralization axis (PC1) to test the classical hypothesis that critical dynamics are governed by central network hubs (Figure 6D). In this 1D framework, physically constrained environments (e.g., 60-module Cortical devices) exhibited moderate coupling (*r* = 0.474). PDMS micro-channels force the creation of structural bottlenecks, ensuring that network dynamics are heavily gated by these physical choke points. Conversely, unconstrained networks, such as Monolayer Cortical (*r* = −0.052) and Organoid Cortical (*r* = 0.182) cultures, appeared entirely uncoupled. Because these open architectures lack physical walls, judging them solely by their reliance on central hubs was a severe mischaracterization of their topological constraints. Therefore, without these physical choke points (PC1) to dictate flow, we hypothesized that their dynamics must be constrained by the secondary topological dimension, spontaneous functional segregation (PC2). To test this, we expanded our analysis to the full 2D state-space using Canonical Correlation Analysis (CCA) (Figure 6E). This multidimensional mapping revealed a profound “Hidden Coupling” phenomenon in Monolayer Cortical networks. While they appeared entirely uncoupled along the PC1 axis, integrating PC2 caused their structure-function coupling to skyrocket to a near-perfect mathematical alignment (CCA = 0.992). Biologically, this suggests that unconstrained 2D monolayers are not devoid of structural control; rather, they control their massive activity by spontaneously self-organizing into isolated functional modules. This establishes a core principle of *in vitro* computational design - that functional segregation (PC2) can substitute perfectly for physical walls (PC1) to achieve deterministic control over network dynamics.

In stark contrast, the inclusion of the full 2D state-space confirmed a true biological decoupling within 3D architectures. Organoid Cortical cultures exhibited weak coupling in 1D (*r* = 0.182) and remained heavily decoupled even when accounting for macroscopic functional modularity (CCA = 0.247). This confirms that unlike flat monolayers, organoids possess true spatial degrees of freedom. Their critical dynamics operate independently of their macroscopic topological blueprint, likely utilizing dense, localized Z-axis microcircuits to route activity; a volumetric strategy that bypasses the rigid 2D macro-constraints entirely [48].

Finally, our analysis revealed that cellular composition acts as a secondary structural constraint, capable of overriding a network’s baseline routing strategy. Across both 2D and 3D environments, the use of highly excitable hippocampal neurons significantly increased the network’s reliance on primary structural hubs. For example, moving to hippocampal cultures from Monolayer Cortical cultures jumped their 1D coupling from *r* = −0.052 up to *r* = 0.616, while Organoid cultures saw a similar jump from *r* = 0.182 to *r* = 0.532. We propose a hippocampal “Anchor” effect: introducing hyper-excitable, burst-prone cell types forces the network to route traffic through its physical choke points (PC1) to manage massive waves of synchronized activity, effectively anchoring the network’s dynamics to its rigid physical infrastructure regardless of its baseline spatial freedom.

## 3 Discussion

The development of SBI hinges on the ability to engineer reproducible computational states within a biological substrate[49]. This challenge requires hardware that supports real-time electrophysiological paradigms and control of stimulation artifacts, software that allows for accessible task development and temporally precise stimulation, and wetware (cell cultures) that is controllable and reproducible[50]. Here we used a new platform, the CL1, to perform RC on a spatio-temporal MNIST classification task [38]. The previously described CL API interface allowed this task to be developed rapidly and with an incredibly high degree of electrophysiological control, including precise sub-millisecond timing and latency [39]. Rather than treating the decoding pipeline as a fixed implementation detail, we used the flexibility and scalability of the CL1 system to systematically evaluate how cellular composition, architectural governance, and decoding methodology interact to dictate neurocomputational capacity.

### Cell type and architecture has a significant influence of electrophysiological network dynamics

The use of a range of cell types and culture architecture in this study provides a novel opportunity to compare how cell types (in this case Hippocampal versus cortical) and culture architecture, (monolayer, organoids, and modular configurations), give rise to distinct electrophysiological characteristics. Specifically, we observed that hippocampal cultures were generally more active than their cortical counterparts. Likewise, there were also distinct differences between the neuro-computationally relevant properties, such as in criticality metrics during spontaneous activity. Previous work has already identified that criticality arises far more strongly when cultures are in a structured information environment [51]. While characterizing these differences elucidates criticality dynamics during spontaneous activity, future investigations must employ closed-loop paradigms to determine how these metrics modulate in response to structured information. Additionally, although primary electrophysiological features (mean, variance, and maximum firing rates) were highly correlated, the mean and variance of the ISI demonstrated notably weaker correlations. This decoupling indicates the presence of heterogeneous activity patterns, specifically highlighting the mechanistic differences between bursting and continuous firing regimes.

### Cellular Composition and the Spatio-Temporal Reservoir

Our results highlight that cellular identity is a foundational driver of information processing. As demonstrated by electrophysiological and functional connectivity analyses, hippocampal lineages exhibited heightened overall activity, highly regular firing patterns, and strong local segregation compared to purely cortical cultures. While hippocampal neurons naturally facilitate the longer temporal traces required to process sequential data, our findings suggest that the computational advantage provided by these lineages is not static; it is heavily mediated by the structural environment in which they are cultured. The ability of iPSC differentiation protocols to generate distinct, region-specific subtypes allowed us to observe that cortical and hippocampal cells, while sharing basic electrophysiological properties, respond to structured spatio-temporal inputs with differing dynamical signatures. Further research is required to consider the functional synergy of co-cultures (e.g. cortical and hippocampal), as the interplay between these two regions is a known key driver of adaptive modeling [41, 52].

### Controlling Cell Culture Structures Highlights the Potential of Enforced Modularity

The variability of biological neural networks (BNNs) is one feature which may prevent reliability and replicability of results. While previous works have explored RC approaches with neural cell cultures in monolayers [5, 53, 54], organoids [7], and in modular setups [21, 55], different setups and methods have precluded a direct comparison and this has previously prevented understanding what structural features, if any, lead to better performance in classification tasks. Here we found strong support that for human iPSC-derived networks, physical cell culture structure (the geometry and topology) is one defining constraint on neurocomputational capacity.

Building on Sumi et al. [21] and Sono et al. [55], who demonstrated that modularity enhances dynamical separability in rat cortical neurons, we showed that this principle holds true in more complex human BNNs. By consistently achieving the highest average accuracies on the spatio-temporal MNIST task using modular cortical cultures across every decoding mechanism, compared to lower performance in unstructured monolayers, organoids, and ~10% in shuffled controls, we provide evidence that engineered modularity significantly outperforms random connectivity for high-order processing. The performance gap between modular networks and unstructured configurations may be attributed to the mechanisms of functional specialization and perturbation isolation highlighted in complex network theory [26, 56]. Unstructured monolayers are often characterized by network-wide bursting that dominates their spontaneous activity [57]. This global synchronization may collapse the high-dimensional state space necessary for complex RC, as the system’s fading memory is periodically reset by these network-wide events, violating the separation property required for complex processing [58].

In contrast, our PDMS 60-module devices enforced a topology that mimics the brain’s small-world architecture [59] at a micro-scale, clustering connections locally while maintaining sparse long-range links. This structure appears to act as a functional regularizer, allowing individual modules to operate semi-independently without propagating perturbations to the entire network. This isolation likely preserved the distinct dynamical trajectories required to separate the complex spatio-temporal patterns of MNIST, effectively balancing the reservoir’s need for integration with the necessity of segregation. Our evaluation using this spatio-temporal task reveals that higher-order temporal processing is deeply topology-dependent. It is highly probable that signal delays and reverberations introduced by microfluidic bottlenecks in the modular system created a richer “fading memory,” allowing the reservoir to capture the time-dependent features encoded by our autoencoder [25, 60]. Ultimately, this implies that while unstructured neural cell culture systems can map static inputs, continuous neurocomputing and complex temporal encoding relies strongly on geometric constraints to restrict the spontaneous dynamics that disrupt stable data representations.

Our spike-count decoding pipeline, which employs trial-based holdouts and strict stimulation-window contamination controls, yields only a modest margin above chance. The methodology used to obtain this result is broadly consistent with recent work by Ciampi et al. [61], who reported a reliable biological baseline of 36% accuracy on the MNIST task using unstructured cortical networks with rigorous artifact removal and randomization. Notably, our observed accuracies for both Time-bin and Frequency-domain mechanisms in comparable monolayer cultures bracket this 36% baseline, suggesting a consistent limit to computational performance in these systems. These performance variations suggest that, like AI architectures, biological substrates may possess specific decoding strengths. Our data indicates that frequency-domain decoding better captures the latent dynamics of these cultures than time-domain metrics, unless time-based holdout is implemented (however the latter likely reflects a different type of information processing, such as fading memory, rather than classification). This implies that decoding performance reflects a functional alignment between the chosen strategy and the network’s intrinsic temporal signatures, rather than a monolithic measure of computational capacity. This convergence, despite differences in implementation, offers validation of our qualitative conclusion: once artifact and cross-validation leakage are properly controlled, biological RC accuracy on this task is well above chance, but considerably more modest than values obtained under less rigorously validated pipelines.

### Rigorous Controls Are Necessary to Mitigate Inflated Accuracies

RC is a potentially powerful approach to use a substrate to transform information. However, using RC to determine whether dynamical systems engage in consistent transformations of information is sensitive to two potential false positive sources. Firstly, a critical challenge is the confounding presence of stimulation artifacts [35]. If these transients are not meticulously managed, machine learning classifiers may easily overfit to the geometric distinctness of the electrical noise rather than decoding true non-linear biological computation, especially if proper controls are not implemented as comparable baselines.

The second source of potential concern is that neural activity can be extremely high-dimensional. While this is an attraction of biological systems, when dimensionality becomes exceptionally large relative to the amount of data, three key issues can arise:

1. Random separability increases, as even randomly arranged data points can be linearly separated in high-dimensional space [62].
2. Overfitting may occur if reservoir dimensions are too large relative to training samples, where even small systematic biases are identified as key features [63].
3. A large feature space also increases the risk of identifying spurious predictive relationships due to multiple testing, feature selection bias, or chance correlations, particularly when the number of features greatly exceeds the number of training samples [64].

Therefore, it is critically important that controls for RC approaches take into account the dimensionality of the dataset and ensure the decoder is unbiased. Our direct comparison of decoding mechanisms illustrates these sources concretely. Our multi-layered framework, including hardware blanking, software-based spike sorting, and shared controls (shuffled data, saline, and synthetic random spike trains), revealed that not all decoding mechanisms are equally trustworthy. The saline control’s elevated accuracy under the frequency-domain with stim mechanism (mean 44.4%) demonstrates that a mechanism can appear to exceed chance while reflecting hardware artifacts; only the combination of a stimulus-excluding response window and a saline control makes this detectable.

It is worth noting that if the primary objective is purely boosting overall classification performance including and exploiting these stimulus artifacts could theoretically be useful. In such hybrid architectures, the geometric distinctness of such stim artifacts might serve as an additional high-dimensional feature to drive up accuracy. This was particularly evident in the ≈ 20% boost in accuracy when comparing Frequency-decoding vs Frequency with Stim decoding conditions (Figure 5C). However, when the objective is to measure the actual computational capabilities of the biological substrate and advance true neurocomputing frameworks, this approach may be fundamentally flawed. Relying on artifact-driven signal boosting obscures the intrinsic, non-linear processing of the living network, making strict artifact mitigation non-negotiable for evaluating cellular performance. Therefore, previous studies without these controls or with controls that only look at reducing the activity (and therefore the dimensionality) of the neural cultures, should also be treated with caution.

### Decoding Methodology: The Unseen Variable

This work demonstrates that the decoding pipeline itself is a first-class experimental variable. Crucially, the validity of this variable relies on our understanding of fading memory, which is the fundamental reservoir computing principle stating that recent inputs exert a greater influence on the current internal state than past inputs [30, 58]. Because our standard decoding involves fitting and freezing a classifier at the start of an experiment, the resulting predictions across trials may be subject to significant temporal separation. This separation can span several hours and encompass hundreds of intermediate inputs between the state during classifier fitting and the state during prediction. This creates a conceptual gap regarding whether current decoding pipelines accurately reflect the true computational capacity of biological neurons operating as reservoir systems.

To investigate this, we evaluated an alternative paradigm by switching from trial-based to time-bin-based holdout, which allows classifiers to utilize information from the most recent input stimulus. We observed that this shift resulted in a significant performance increase, specifically a more than two-fold improvement in average accuracy and a three-fold increase in variability, which supports the existence of biological fading memory. While time-bin-based holdout is frequently criticized for the potential for data leakage that can inflate performance [65], we observe that the saline and synthetic (SDK) controls do not exhibit this performance boost. The absence of this effect in non-biological controls suggests that the observed increase in biological networks is not a simple statistical artifact but rather an indication of emergent temporal dynamics. The divergence between the biological networks, which show clear improvement under time-bin decoding, and the saline/in-silico controls, which remain at the chance level regardless of the holdout strategy, suggests that the fading memory driving these results could be a genuine property of the biological substrate. While we acknowledge that time-bin-based holdout can yield overconfident conclusions, particularly when used without comparative controls, the observed results suggest that current methodologies do not capture the full extent of the reservoir’s dynamics. We posit that the true predictive performance likely lies between these two paradigms. For example, Hippocampal variants displayed a hyper-deviated, structurally loose state that might have led to a sustained stimulus-independent reverberation with a fading memory dynamic. This very well may have accounted for their high decoding accuracies under specific decoding regimes. Whether this is a useful computational property or a more trivial aspect of this cell culture that happens to lead to higher performance under certain decoding regimes is an open question and depends more on what the research question is or the intended use case than anything else. The key takeaway from this finding is not that these cultures are incapable of displaying some useful RC properties, but that it is vital to be careful of interpretations and assumptions.

We also demonstrate that frequency-domain (PSD) decoding, which utilizes the power spectral density of the signal, serves as a robust alternative to time-binning. While time-binning is highly sensitive to the precise temporal alignment and autocorrelation of spike events, the frequency-domain approach characterizes the oscillatory state of the network rather than the instantaneous spike count. This approach effectively eliminates the influence of temporal jitter.

The presence of fading memory in biological substrates has significant implications for applied biocomputing. Because the biological substrate naturally integrates historical context into its present state without requiring the discrete and energy-intensive memory buffers used in digital architectures, it is inherently optimized for processing sequential and temporal data. This intrinsic autocorrelation could be leveraged to solve complex, time-dependent problems such as real-time signal processing, continuous motion prediction, or dynamic temporal forecasting, offering a uniquely energy-efficient approach to temporal integration.

Finally, the non-stationary nature of living neural cultures necessitates the consideration of how decoding models evolve alongside the network. Because biological networks are continuously adaptive and do not remain static, we expect that future research will require the development of adaptive decoding algorithms. Recent developments in adaptive machine learning algorithms, such as Test-Time-Training (TTT) [66], may inspire new decoding frameworks to characterize the performance of continuously evolving, non-stationary biological reservoirs. For instance, in this paper we observed that monolayers appear to self-organize into isolated functional modules. Previously, the presence of such functional modules has been observed to be vital in order to obtain higher performance in closed-loop learning paradigms [67, 68].

### Limitations and Future Directions

One potential limitation of the current study was inconsistent and, at times, small group sizes due to availability of cultures. In particular, the group sizes of hippocampal cultures were smaller than cortical cultures. The decision to retain these cultures were made as effect sizes were typically large and important differences could still be reliably inferred based on the results. However, future work will benefit from ensuring more balanced group sizes in future studies to allow more nuanced analysis to be conducted equally between groups. Likewise, this current study only looked at cortical cultures and hippocampal cultures. It did not systematically explore the range of other important cell types that exist in neural systems nor how different combinations of them might interact. Future work should explore whether performance can be improved through more complex neural cultures enriched with a variety of cell types. Finally, it must also be recognised that these results do not reflect single trial performance, but a decoding of global responses following repeated presentations of the chosen dataset. Therefore, another limitation is that as this study focused only on RC, there was no closed-loop feedback aiming to train neural cultures. As such, the results reflect the capacity of different decoders being applied to a range of neural cultures, but do not reflect the capacity for these neural systems to learn to respond in a single trial. Future work should aim to explore not only whether neural cultures display characteristics that allow for RC-type performance, but whether this performance can be improved over time with specific feedback following each individual trial.

## 4 Conclusion

This study demonstrates that the computational performance of SBI is not merely a product of the biological substrate’s inherent dynamics, but a complex intersection of cellular identity, architectural topology, and the methodological rigor of the decoding pipeline. We suggest that network structure, particularly through modular PDMS scaffolds, acts as a vital functional regularizer that prevents signal saturation typical of unstructured networks and enables the high-dimensional state space required for complex spatio-temporal tasks like MNIST. Crucially, our results also reveal that classification accuracy is highly sensitive to temporal separation within the decoding framework. While stringent trial-based cross-validation is necessary to distinguish genuine neurocomputational capacity from statistical leakage, our comparison with time-bin-based holdouts indicates that the true predictive potential of the biological reservoir, driven by its intrinsic fading memory, likely lies between these two paradigms. By demonstrating that frequency-domain decoding effectively mitigates temporal autocorrelation pitfalls while preserving this continuous temporal integration, we provide a validated blueprint for evaluating SBI systems. Furthermore, because living neural cultures are inherently non-stationary, successfully leveraging their energy-efficient capacity for time-dependent processing will necessitate a shift toward dynamic, adaptive decoding algorithms. While these results did significantly support the use of modular cultures under this experimental paradigm, this does not indicate that other culture types, such as organoids, are not suitable for neurocomputing tasks. However, results do indicate that care in the interactions for encoding and decoding, including how to stimulate and record from these cells, is vital. Moving forward, these insights into the need for architectural controls and adaptive analytical frameworks will be essential for the reproducible scaling of biocomputing, ensuring that future advancements are built upon a reliable, validated foundation of neurocomputational potential.

## 5 Methods

### 5.1 Morphology and Organization of Neural Cell Cultures

In vitro cell generation is used to advance our understanding of in vivo cell behavior and fundamental biophysical environments which leads to the formation and function of the tissue or organ. Deciphering these processes is key to understanding the complicated organization of these cells which can then be closely mimicked and studied in vitro. Neural cultures can be grown in both flattened structures mainly termed as monolayers, organoids and more recently the above morphologies in microfluidic based systems. Predominately conventional monolayer cultures have been widely used to study neural differentiation, however organoid cultures are shown to more closely mimic the in vivo environment and provide improved neural differentiation and function [69]. The recent advances in neural engineering have led to the development of microfluidic devices which are shown to provide a targeted control of cell growth including equal distribution in spatial and temporal organization of the cells [69]. These microstructures have been designed and fabricated to culture neural tissues with the possibility of understanding the advanced neural migration and provide insights into how these physical cues influence neural plasticity and learning.

Exploring the neural behavior of such lineage specific neurons in various morphologies enables us to explore the information processing and learning behavior of these in-vitro neurons providing a viable direction in the future research of biological computation.

### 5.2 Differentiation of iPSCs to Cortical and Hippocampal Neurons

With the advancement in the field of stem cell differentiation, it is now possible to efficiently differentiate aciPSCs into lineage specific neurons exhibiting spontaneous activity [70]. Targeted region-specific differentiation into cortical and hippocampal neurons was achieved following established protocols with minor optimizations [12, 71, 72].

#### 5.2.1 Data Selection Criteria

All recordings with less than 500 spikes during the recording were excluded for this study (see Supplementary Materials for full dataset). Furthermore, any recordings where potential stimulation artifacts were seen were either removed, or the stimulation artifacts were manually removed before analysis via spike-sorting, described in 5.3.

#### 5.2.2 Monolayer

To test the electrophysiology of the neural networks, MEA chips were used to culture these neurons. MEA chips (Multi Channel Systems) were coated with poly-D-Lysine (PDL) and human Laminin-521 (Stem-Cell Technologies) at a concentration of 50 µg*/*mL and 10 µg*/*mL, respectively. Monolayer cortical neurons (day in vitro (DIV) 30) and hippocampal neurons (DIV 40) were dissociated and seeded either alone (referred to in results as “cortical”, or “hippo”) at 0.5 × 10^6^ total cells per MEA chip in basal differentiation medium, BDM supplemented with 50 nM Chroman 1. Basal differentiation medium consisted of 1:1 DMEM/F12 and Neurobasal with 0.5× B27, 0.5× N2, 0.5× ITSA, 1× GlutaMAX, 0.5× penicillin/streptomycin, and 50 µM 2-mercaptoethanol (Life Technologies). After 24h, Chroman 1 was withdrawn and cells were maintained in basal differentiation medium supplemented with Ara-C (Cytosine arabinoside). Cells were maintained in BrainPhys maturation medium supplemented with brain-derived neurotrophic factor 10 ng*/*mL (BDNF, R&D Systems), 0.25 µM N6,2’-O-dibutyryladenosine 3’,5’-cyclic monophosphate (dbcAMP, Tocris Bioscience), 200 nM ascorbic acid (Sigma-Aldrich) and 10 µM *γ*-secretase inhibitor After 7 days DAPT was removed, and cells were maintained in BrainPhys Neuronal Medium and SM1 Kit (Stem-Cell Technologies) supplemented with BDNF, dbcAMP and ascorbic acid for further testing and analysis. RC testing began on DIV 60.

#### 5.2.3 PDMS Film 60-module Device

Neuronal cultures were seeded at DIV 58 for Monolayers and DIV 70-80 for Organoids into a microfluidic PDMS devices designed to enforce modular network topology. The device features an array of 60 octagonal microfluidic wells, each 150 µm in diameter, connected to eight adjacent wells via microchannels (see Figure 2A). The width and height of the microchannels were 7.1 ± 0.18 µm and 4.4 ± 0.31 µm, respectively, as measured by laser confocal microscopy (Keyence VK-X260; mean ± SD, *n* = 10 microchannels). Crucially, the geometry aligns such that each well overlays a single electrode node. This PDMS architecture facilitates the development of a highly modular network, in contrast to unconstrained monolayer cultures where neurons form connections without spatial restriction. The structures were designed in Autodesk Fusion 360 and fabricated via soft lithography, yielding a PDMS film thickness of approximately 100 µm [73, 74].

Following film preparation and sterilization, the PDMS microfluidic films were aligned onto the electrode area of the MEA chips with each well overlaying a single electrode node (The MEA chips used for this study have 64 electrodes). To allow maximum adhesion to the chip, the MEAs were left overnight in the biosafety cabinet to promote drying. The following day, cortical or hippocampal neurons were dissociated and seeded onto these MEAs as described in the monolayer subsection. It should be noted that PDMS films did not create a perfect constraint on neural connectivity, as neural growth continued under the PDMS wells to some degree. It is likely this undirected growth allowed for some isolated long range connections between otherwise segregated communities. While this was unintended, it is interested to note that this means these cultures had a balance between containing cells within a highly segregated octagonal well-like structures and allowing for long-range axonal connections between these structures. The resulting functional connectivity mimics the “small-world” properties, inherent to many brain structures [59]. RC testing began on DIV 60.

#### 5.2.4 Organoid

Neural cultures were differentiated from iPSCs according to a previously published protocol [12] until DIV 19, at which point cells were cryopreserved. Briefly, DIV 19 neuro precursor cells (NPCs) were thawed and seeded at 0.5 × 10^6^ cells*/*cm^2^ in BDM supplemented with 50 nM Chroman 1. After 24 h, Chroman 1 was withdrawn and cells were maintained in BDM for 2 d. The medium was then supplemented with 3 µM CHIR99021 (Wnt activator) till DIV 35. To generate hippocampal organoids (HOs), cells were seeded at 50 000 to 60 000 per well of a 96 ultra-low attachment multiple well plate (Sigma-Aldrich CLS7007) in CTX media with 25 nM Chroman 1. After 48 h, Chroman 1 was removed and cells were maintained in BDM for 14 d with media change every other day. The organoids were transferred into a 10 cm Petri dish with maturation medium supplemented with 10 ng*/*mL BDNF, 0.25 µM dbcAMP, 200 nM ascorbic acid, and cultured on an orbital shaker (Biotek #NBT101SRC) in the incubator at 68 rpm.

To generate cortical organoids (CO), neural cultures were differentiated from iPSCs according to previously published protocol [12, 71, 72] until DIV 19. DIV 19 NPCs were seeded at 50 000 to 60 000 per well of a 96 ultra-low attachment multiple well plate (Sigma-Aldrich CLS7007) in BDM media with 25 nM Chroman 1. After 48 h, Chroman 1 was removed and cells were maintained in BDM media until DIV 30 with media change every other other day. The organoids were transferred into a 10 cm Petri dish and cultured as the HOs as above.

For electrophysiological studies, HOs were transferred onto PDL/laminin-521-coated MEA chips (Multi Channel Systems). MEA chips were coated with PDL and human Laminin-521 at a concentration of 50 µg*/*mL and 10 µg*/*mL, respectively. The organoids at DIV 55 were transferred onto PDL/laminin-521-coated chips. After 48 hours, Chroman 1 was removed and cultured in Brainphys medium supplemented with 10 ng*/*mL BDNF, 0.25 µM dbCAMP, and 200 nM ascorbic acid. RC testing began on DIV 120.

### 5.3 Stimulation and Spike Sorting

Each stimulation in this study was a negative-leading biphasic pulse (see Figure 7). The stimulation current and pulse width were set at 2 µA and 160 µs per phase, respectively, for all trials. Recording is blanked for 2ms after stimulation to reduce artifacts.

**Fig. 7.**
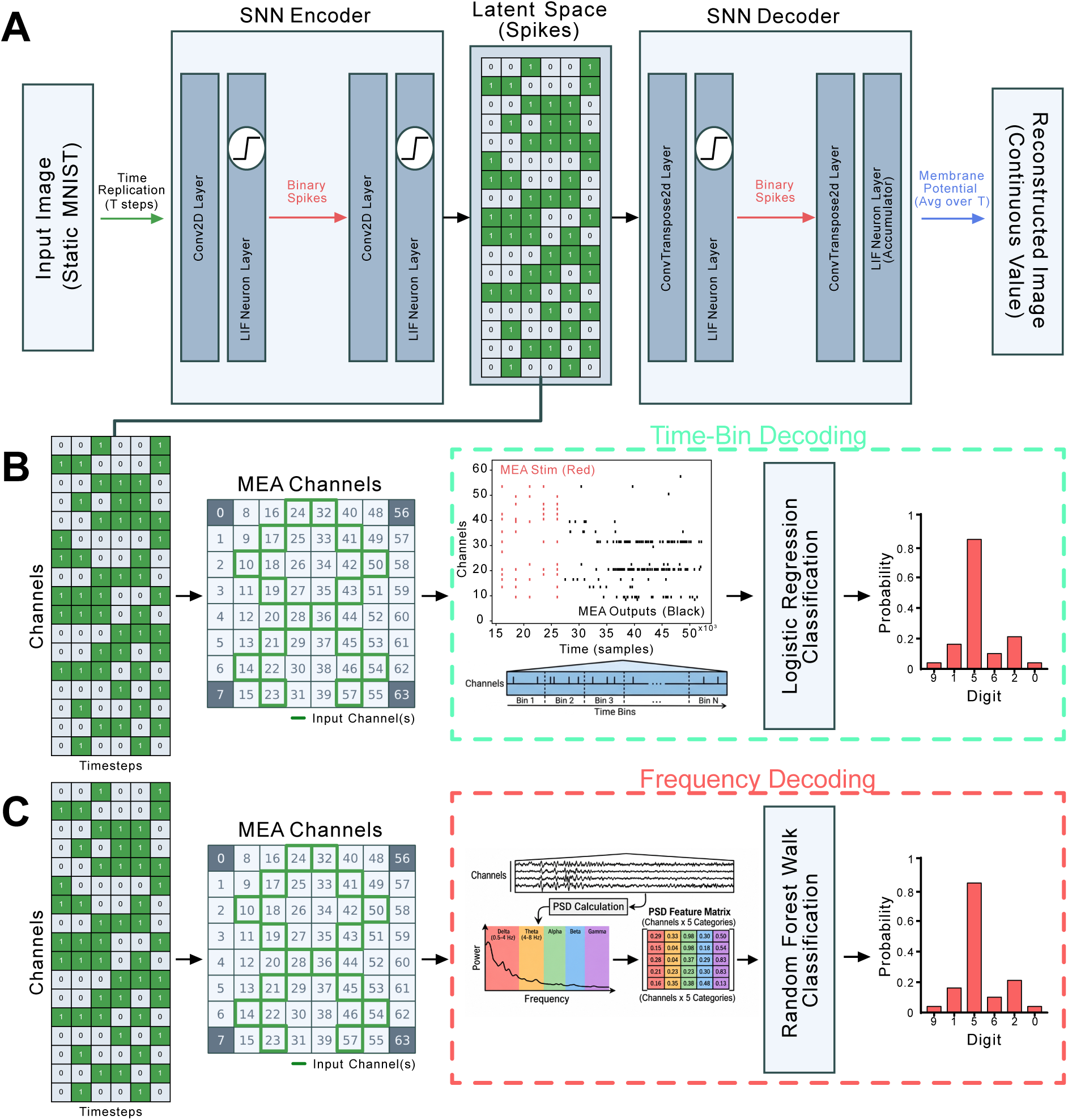
Pre-processing of MNIST image and stimulation protocol. **A:** A static MNIST image is converted to a spatiotemporal spiking signal using a spiking autoencoder. The static MNIST image is flattened and replicated *T* times to convert it into a temporal signal. The resulting tensor is reduced into a smaller spatio-temporal signal of *N* by *t* via the autoencoding process. In this example *N* = 16 and *t* = 5. After training the autoencoder, the spatio-temporal latent signal is extracted and fed in as input to 16 channels of the MEA over 5 steps, delivered as a series of biphasic stimulation pulses. The resulting neural response was then decoded using each of the four mechanisms described in Decoding Mechanism 3 & 4: Time-Binned Spike-Count Decoding, Trial Holdout vs. Time Holdout (**B**) and Decoding Mechanism 1 & 2: Frequency-Domain (PSD) Decoding, Excluding vs. Including the Stimulation Window (**C**).

A one second recording was taken after the last stimulation of each trial. During this time, spikes were detected in real-time at a sampling rate of 25 kHz, and saved to HDF5 recordings. A 75-sample waveform cutout around each detected spike (3ms total; 1 ms prior and 2 ms after) was also saved in the file. Waveforms for each file were collated, sorted and clustered using Principle Component Analysis (PCA) and Gaussian Mixture Models (GMM). Clusters that were deemed noisy or invalid (e.g. stimulation artifacts) by two independent experts were removed, and RC was then recomputed with a reduced waveform pool. This method was employed to ensure the classification accuracy was not influenced by stimulation or noise artifacts.

### 5.4 Classification

To test spatio-temporal performance of neurons, we performed the classic MNIST handwritten digit recognition task on different types of neuronal cultures (Monolayer, 60-module and Organoids). A total of 23 unique cell cultures were tested over 50 recordings (after excluding non-viable cultures; see Supp Tables S1, S2, S3, S4, and S5 for all decoding results).

#### Conversion of Static Image to Spatio-temporal Spike-train

For this task, 60,000 digits were converted from a static 28 × 28 pixel image to a spatial-temporal spike matrix of *C* input channels by *T* timesteps (see Figure 7). A fully-spiking autoencoder was used to convert the pixels of a hand-written digit into a spatio-temporal matrix. Both the encoder and decoder were 3 layer fully-connected CNNs with LIF neurons. The autoencoder was inspired by a fully-spiking variation autoencoder introduced by Kamata et al. [75]. However, the fully spiking CNNs and LIF models used in this study were implemented via the snnTorch package [47]. Briefly, static images were flattened into a 1D 784-element vector, and then replicated *T* = 5 times to create a 2D spatio-temporal matrix. They are fed into the autoencoder, which acts as both a spatio-temporal feature extractor and a dynamic compression mechanism [75]. During the forward pass, the integrate-and-fire dynamics of the LIF neurons encode the static spatial features into discrete spike events distributed across the *T* simulated timesteps. The encoder condenses the flattened inputs into a lower-dimensional (16 × 5) latent spike representation, while the decoder reconstructs the original static image from these dynamic spike trains. Because the thresholding function of a spiking neuron is non-differentiable, the network was trained using backpropagation through time (BPTT) coupled with a surrogate gradient approach. By minimizing the reconstruction loss (MSE) the autoencoder learns to generate a rich, sparse spatio-temporal representation of the digits. Once trained, the spike tensor extracted from the latent layer serves as the optimal dynamic input for the downstream components of the model.

We fed a subset of 1000 spatio-temporal spike-train representations of digits into MEA electrodes one after the other (see Figure 7). For all results presented here, 16 input channels and 5 timesteps were used to encode each digit stimulus. Each timestep was separated by 100 ms, with stimulation occurring at a frequency of 10 Hz. From each such recording, we then applied the four decoding mechanisms offline as described below, so that all comparisons in this manuscript are drawn from identical underlying stimulation and recording sessions.

#### Decoding Mechanism 1 & 2: Frequency-Domain (PSD) Decoding, Excluding vs. Including the Stimulation Window

The neurons’ response was operationalised as the raw read-out voltage on each of 59 electrodes. For the Frequency-domain mechanism, this was recorded for a 400 ms window beginning 10 ms after the final stimulation pulse of each trial; this delay was chosen to exclude residual stimulation artifact, consistent with our blanking approach described in section 5.3. Blanking was increased in software to 10ms to ensure no residual lingering of artifacts. For the Frequency-domain with Stim mechanism, the identical downstream feature extraction and classifier were applied, but the window began at the onset of stimulation rather than 10 ms after its offset, so that the response necessarily includes the stimulation period itself. In both cases, the window was segmented into consecutive 10 ms bins, and for each bin the power spectral density (PSD) of the raw signal on each channel was estimated via a Fast Fourier Transform (FFT). PSD values were then aggregated into five canonical frequency bands (delta, theta, alpha, beta, and gamma), yielding a five-band feature vector per channel per trial. This feature representation, *X*, was used to train a Random Forest classifier to predict the digit label *y* (0–9) for each trial, with 20% of trials held out per fold (cross-trial holdout).

#### Decoding Mechanism 3 & 4: Time-Binned Spike-Count Decoding, Trial Holdout vs. Time Holdout

For comparison to the frequency-domain mechanisms above, we also decoded the same underlying recordings using the original time-binned spike-count approach. Spike counts were discretised into temporal bins and summed across trials to produce a feature tensor *X* with dimensions (*N* × *C* × *T* ), where *N* is the number of trials, *C* the number of channels (59), and *T* the number of time bins. This tensor was flattened to two dimensions (*N* , *C* × *T* ) and used to train a logistic regression classifier to predict digit identity. For the Time Bin (Trial Holdout) mechanism, cross-validation held out 20% of trials (the *N* -axis) per fold, consistent with the cross-trial holdout used for the frequency-domain mechanisms above. For the Time Bin (Time Holdout) mechanism, the identical feature tensor and classifier were used, but cross-validation instead held out time bins (the *T* -axis) rather than trials.

#### Average Probability Decoding

To mitigate the high trial-to-trial variability inherent in living neural networks, we utilized an average probability decoding approach for all main reported results (Figure 5C&D). Because single-trial predictions can be heavily influenced by spontaneous biological noise, we computed a prediction by aggregating output probabilities. Specifically, for each target trial, the classifier’s predicted probability distributions were averaged alongside four additional trials of the same stimulus class. The digit class with the highest mean probability across this five-trial ensemble was taken as the final prediction. Baseline single-trial classification accuracies are detailed in Supp Table S5.

#### 5.4.1 Controls

Three shared control conditions were applied identically across all four decoding mechanisms described above. First, a shuffled control was constructed by randomly shuffling spike counts (or, for the frequency-domain mechanisms, the corresponding raw-signal segments) across channels and time bins within each recording, to create a fully randomised dataset that retained an identical culture-wide firing rate (see Supp Figure S3). Second, a saline (no-cell) control was recorded by delivering identical stimulation into a saline-only dish (without neuronal cells), and decoding it with each of the four mechanisms in turn; this isolates any component of apparent accuracy attributable to hardware or electrical artifact rather than genuine neuronal activity. Third, a synthetic random-spike control was generated independently of any recorded neural or saline data (i.e., not exposed to real stimulation events), and decoded with each of the four mechanisms in turn, to rule out systematic bias in the encoding or decoding software itself. An additional control method was implemented using a test-chip instead of an MEA cell culture - a simple RC circuit designed to mimic the electrical properties of saline solution, typically used to calibrate the MEA, which acts as a further control for non-cellular electrical activity.

#### 5.4.2 Activity, Criticality and Functional Connectivity Metrics

PCA was selected as the primary dimensionality reduction technique to construct Functional Connectivity (FC), Activity, and Criticality indices. The use of PCA was necessitated by three factors inherent to MEA data. First, it resolves multi-collinearity; raw network metrics (e.g., mean firing rate and maximum spike count) are often highly inter-correlated, which can severely confound downstream linear models and clustering algorithms [76, 77]. PCA optimally weights and combines these redundant features into singular axes of variance, a standard approach for distilling high-dimensional, collinear electrophysiological data [78, 79].

Second, it extracts latent macroscopic states. In complex neural systems, the overarching dynamical regime (critical state) is a distributed network property rather than a single measurable variable [80, 81]. By evaluating the first and second principal components (PC1 and PC2), we capture the dominant axes of this systemic variance. Finally, PCA generates strictly orthogonal components [76]. This ensures that the inputs fed into our subsequent Canonical Correlation Analysis (CCA) algorithms represent mathematically independent feature spaces, thereby preventing the artificial inflation of correlation scores during multidimensional moderation analysis.

To determine how number of active channels affected FC and accuracy, active channels were defined as the number of channels out of 60 which had higher than 75% percentile of spiking activity (i.e. on average around 1400 spikes per recording).

## Acknowledgments

This work is supported by funding from the Australian Government’s Industry Growth Program and by the U.S. Defense Advanced Research Projects Agency (DARPA). The views, opinions, and/or findings expressed are those of the author(s) and should not be interpreted as representing the official views or policies of the Department of Defense or the U.S. Government.

## Declarations

The authors declared the following potential conflicts of interest with respect to the research, authorship, and/or publication of this article: B.J.K., F.H., B.W., A.A., A.L., J. Z., C.D., K.D.A., H.W.C. and F.D. are employees of and may hold shares or another interest in Cortical Labs, a research-focused start-up working in a space and holding patents related to this article. No specific incentive was provided to any author for contribution to this article.

## References

[1] Smirnova, L.: Biocomputing with organoid intelligence. Nature Reviews Bioengineering 2(8), 633–634 (2024) 10.1038/s44222-024-00200-6

[2] Kagan, B.J.: Two roads diverged: Pathways toward harnessing intelligence in neural cell cultures. Cell Biomaterials 1(8), 100156 (2025) 10.1016/j.celbio.2025.100156

[3] Shahaf, G., Marom, S.: Learning in Networks of Cortical Neurons. The Journal of Neuroscience 21(22), 8782–8788 (2001) 10.1523/JNEUROSCI.21-22-08782.2001

[4] Bakkum, D.J., Chao, Z.C., Potter, S.M.: Spatio-temporal electrical stimuli shape behavior of an embodied cortical network in a goal-directed learning task. Journal of Neural Engineering 5(3), 310–323 (2008) 10.1088/1741-2560/5/3/004

[5] Isomura, T., Kotani, K., Jimbo, Y.: Cultured Cortical Neurons Can Perform Blind Source Separation According to the Free-Energy Principle. PLOS Computational Biology 11(12), 1004643 (2015) 10.1371/journal.pcbi.1004643

[6] Kagan, B.J., Kitchen, A.C., Tran, N.T., Habibollahi, F., Khajehnejad, M., Parker, B.J., Bhat, A., Rollo, B., Razi, A., Friston, K.J.: In vitro neurons learn and exhibit sentience when embodied in a simulated game-world. Neuron 110(23), 3952–39698 (2022) 10.1016/j.neuron.2022.09.001

[7] Cai, H., Ao, Z., Tian, C., Wu, Z., Liu, H., Tchieu, J., Gu, M., Mackie, K., Guo, F.: Brain organoid reservoir computing for artificial intelligence. Nature Electronics 6(12), 1032–1039 (2023) 10.1038/s41928-023-01069-w

[8] Robbins, A., Schweiger, H.E., Hernandez, S., Spaeth, A., Voitiuk, K., Parks, D.F., Van Der Molen, T., Geng, J., Cline, I., Kosik, K.S., Salama, S.R., Sharf, T., Mostajo-Radji, M.A., Haussler, D., Teodorescu, M.: Goal-directed learning in cortical organoids. Cell Reports 45(2), 116984 (2026) 10.1016/j.celrep.2026.116984

[9] Tessadori, J., Bisio, M., Martinoia, S., Chiappalone, M.: Modular Neuronal Assemblies Embodied in a Closed-Loop Environment: Toward Future Integration of Brains and Machines. Frontiers in Neural Circuits 6 (2012) 10.3389/fncir.2012.00099

[10] Pimashkin, A., Gladkov, A., Mukhina, I., Kazantsev, V.: Adaptive enhancement of learning protocol in hippocampal cultured networks grown on multielectrode arrays. Frontiers in Neural Circuits 7 (2013) 10.3389/fncir.2013.00087

[11] Zhang, X., Dou, Z., Kim, S.H., Upadhyay, G., Havert, D., Kang, S., Kazemi, K., Huang, K., Aydin, O., Huang, R., Rahman, S., Ellis-Mohr, A., Noblet, H.A., Lim, K.H., Chung, H.J., Gritton, H.J., Saif, M.T.A., Kong, H.J., Beggs, J.M., Gazzola, M.: Mind In Vitro Platforms: Versatile, Scalable, Robust, and Open Solutions to Interfacing with Living Neurons. Advanced Science 11(11), 2306826 (2024) 10.1002/advs.202306826

[12] Abu-Bonsrah, K.D., Desouza, C., Habibollahi, F., Chan, H.W., Watmuff, B., Dottori, M., Kagan, B.J.: A novel protocol for the efficient generation of all three major hippocampal neuronal sub-populations from human pluripotent stem cells. bioRxiv. ISSN: 2692-8205 Pages: 2026.01.21.700748 Section: New Results (2026). 10.64898/2026.01.21.700748. https://www.biorxiv.org/content/10.64898/2026.01.21.700748v1

[13] Shirzadi, S., Dadgostar, M., Hosseinzadeh, H., Einalou, Z.: Dynamics of frontal cortex functional connectivity during cognitive tasks: insights from fNIRS analysis in the Dual n-back Paradigm. Cognitive Processing 26(3), 555–566 (2025) 10.1007/s10339-025-01275-8

[14] Stone, H.L., Mitchell, J.L., Fuentes-Jimenez, M., Tran, J.E., Yeatman, J.D., Yablonski, M.: Anatomically Distinct Regions in the Inferior Frontal Cortex Are Modulated by Task and Reading Skill. The Journal of Neuroscience 45(19), 1767242025 (2025) 10.1523/JNEUROSCI.1767-24.2025

[15] Chau, B.K.H., Law, C.-K., To, J.Y.L., Shum, D.H.K., Mars, R.B.: Complex functions of human lateral frontopolar cortex. Brain 148(11), 3833–3843 (2025) 10.1093/brain/awaf289

[16] Martin, S., Ferrante, M., Bruera, A., Hartwigsen, G.: Causal contributions of left inferior and medial frontal cortex to semantic and executive control. Communications Biology 8(1), 1343 (2025) 10.1038/s42003-025-08848-5

[17] Bird, C.M., Burgess, N.: The hippocampus and memory: insights from spatial processing. Nature Reviews Neuroscience 9(3), 182–194 (2008) 10.1038/nrn2335

[18] Nakazawa, K., Quirk, M.C., Chitwood, R.A., Watanabe, M., Yeckel, M.F., Sun, L.D., Kato, A., Carr, C.A., Johnston, D., Wilson, M.A., Tonegawa, S.: Requirement for Hippocampal CA3 NMDA Receptors in Associative Memory Recall. Science 297(5579), 211–218 (2002) 10.1126/science.1071795

[19] Paulson, A.L., Zhang, L., Prichard, A.M., Singer, A.C.: 40 Hz sensory stimulation enhances CA3-CA1 coordination and prospective coding during navigation in a mouse model of Alzheimer’s disease. Neuroscience (2024). 10.1101/2024.10.23.619408. http://biorxiv.org/lookup/doi/10.1101/2024.10.23.619408

[20] Gao, Y., Qiao, Y., Wang, X., Zhu, M., Yu, L., Yuan, H., Li, L., Hu, N., Xu, J.-T.: Downregulation of Neuralized1 in the Hippocampal CA1 Through Reducing CPEB3 Ubiquitination Mediates Synaptic Plasticity Impairment and Cognitive Deficits in Neuropathic Pain. Neuroscience Bulletin 41(12), 2233–2253 (2025) 10.1007/s12264-025-01536-8

[21] Sumi, T., Yamamoto, H., Katori, Y., Ito, K., Moriya, S., Konno, T., Sato, S., Hirano-Iwata, A.: Biological neurons act as generalization filters in reservoir computing. Proceedings of the National Academy of Sciences 120(25), 2217008120 (2023) 10.1073/pnas.2217008120. Publisher: Proceedings of the National Academy of Sciences

[22] Rodriguez, N., Izquierdo, E., Ahn, Y.-Y.: Optimal modularity and memory capacity of neural reservoirs. Network Neuroscience 3(2), 551–566 (2019) 10.1162/netna00082

[23] Moriya, S., Yamamoto, H., Hirano-Iwata, A., Kubota, S., Sato, S.: Quantitative Analysis of Dynamical Complexity in Cultured Neuronal Network Models for Reservoir Computing Applications. In: 2019 International Joint Conference on Neural Networks (IJCNN), pp. 1–6 (2019). 10.1109/IJCNN.2019.8852207. ISSN: 2161-4407. https://ieeexplore.ieee.org/document/8852207

[24] Sporns, O., Tononi, G., Edelman, G.M.: Theoretical Neuroanatomy: Relating Anatomical and Functional Connectivity in Graphs and Cortical Connection Matrices. Cerebral Cortex 10(2), 127–141 (2000) 10.1093/cercor/10.2.127

[25] Yamamoto, H., Moriya, S., Ide, K., Hayakawa, T., Akima, H., Sato, S., Kubota, S., Tanii, T., Niwano, M., Teller, S., Soriano, J., Hirano-Iwata, A.: Impact of modular organization on dynamical richness in cortical networks. Science Advances 4(11), 4914 (2018) 10.1126/sciadv.aau4914

[26] Espinosa-Soto, C., Wagner, A.: Specialization Can Drive the Evolution of Modularity. PLOS Computational Biology 6(3), 1000719 (2010) 10.1371/journal.pcbi.1000719. Publisher: Public Library of Science

[27] Yamamoto, H., Spitzner, F.P., Takemuro, T., Buendía, V., Murota, H., Morante, C., Konno, T., Sato, S., Hirano-Iwata, A., Levina, A., Priesemann, V., Muñoz, M.A., Zierenberg, J., Soriano, J.: Modular architecture facilitates noise-driven control of synchrony in neuronal networks. Science Advances 9(34), 1755 (2023) 10.1126/sciadv.ade1755

[28] Duenki, T., Ikeuchi, Y.: Multi-organoid loop cerebral connectoids exhibit enhanced neuronal network dynamics and sequence-specific entrainment. Communications Biology 9(1), 302 (2026) 10.1038/s42003-026-09589-9

[29] Jimbo, Y., Robinson, H.P.C., Kawana, A.: Strengthening of synchronized activity by tetanic stimulation in cortical cultures: application of planar electrode arrays. IEEE Transactions on Biomedical Engineering 45(11), 1297–1304 (1998) 10.1109/10.725326

[30] Jaeger, H.: Short term memory in echo state networks. GMD-Report 152. GMD - GERMAN NATIONAL RESEARCH INSTITUTE FOR COMPUTER SCIENCE (2002)

[31] Tanaka, G., Yamane, T., Hèroux, J.B., Nakane, R., Kanazawa, N., Takeda, S., Numata, H., Nakano, D., Hirose, A.: Recent advances in physical reservoir computing: A review. Neural Networks 115, 100–123 (2019) 10.1016/j.neunet.2019.03.005

[32] Lukoševičius, M., Jaeger, H.: Reservoir computing approaches to recurrent neural network training. Computer Science Review 3(3), 127–149 (2009) 10.1016/j.cosrev.2009.03.005

[33] Tino, P.: Dynamical systems as temporal feature spaces. J. Mach. Learn. Res. 21(1), 44–1649441690 (2020)

[34] Ju, H., Dranias, M.R., Banumurthy, G., VanDongen, A.M.J.: Spatiotemporal Memory Is an Intrinsic Property of Networks of Dissociated Cortical Neurons. Journal of Neuroscience 35(9), 4040–4051 (2015) 10.1523/JNEUROSCI.3793-14.2015. Publisher: Society for Neuroscience Section: Articles

[35] Wagenaar, D.A., Potter, S.M.: Real-time multi-channel stimulus artifact suppression by local curve fitting. Journal of Neuroscience Methods 120(2), 113–120 (2002) 10.1016/S0165-0270(02)00149-8

[36] Gnadt, J.W., Echols, S.D., Yurgelun-Todd, D., et al.: Spectral cancellation of microstimulation artifact for simultaneous neural recording in situ. IEEE Transactions on Biomedical Engineering 50(10), 1129–1139 (2003) 10.1109/TBME.2003.816077

[37] Harrison, R.R., Charles, C.: A low-power low-noise cmos amplifier for neural recording applications. IEEE Journal of Solid-State Circuits 38(6), 958–965 (2003)

[38] Kagan, B.J.: The CL1 as a platform technology to leverage biological neural system functions. Nature Reviews Bioengineering (2025) 10.1038/s44222-025-00340-3. Publisher: Springer Science and Business Media LLC

[39] Hogan, D., Doherty, A., Khoo, B.K., Zhou, J., Salib, R., Stewart, J., Lawson, K., Loeffler, A., Kagan, B.: CL API: Real-Time Closed-Loop Interactions with Biological Neural Networks (2026). https://arxiv.org/abs/2602.11632v1

[40] Zhou, J., Tanneberg, D., Habibollahi, F., Loeffler, A., Lawson, K., Baccetti, V., Abu-Bonsrah, K.D., Desouza, C., Doensen, F., Watmuff, B., Kornienko, D., Azadi, A., Bourke, J.L., Sendhoff, B., Kagan, B.J.: Embodied Neurocomputation: A Framework for Interfacing Biological Neural Cultures with Scaled Task-Driven Validation. arXiv. arXiv:2605.13315 [cs.ET] (2026). 10.48550/arXiv.2605.13315. http://arxiv.org/abs/2605.13315

[41] Sigurdsson, T., Duvarci, S.: Hippocampal-Prefrontal Interactions in Cognition, Behavior and Psychiatric Disease. Frontiers in Systems Neuroscience 9 (2016) 10.3389/fnsys.2015.00190

[42] Rubin, R.D., Watson, P.D., Duff, M.C., Cohen, N.J.: The role of the hippocampus in flexible cognition and social behavior. Frontiers in Human Neuroscience 8 (2014) 10.3389/fnhum.2014.00742

[43] Eichenbaum, H.: Hippocampus: Cognitive Processes and Neural Representations that Underlie Declarative Memory. Neuron 44(1), 109–120 (2004) 10.1016/j.neuron.2004.08.028

[44] Opitz, B.: Memory Function and the Hippocampus. Frontiers of neurology and neuroscience 34, 51–9 (2014) 10.1159/000356422

[45] Rubinov, M., Sporns, O.: Complex network measures of brain connectivity: uses and interpretations. NeuroImage 52(3), 1059–1069 (2010) 10.1016/j.neuroimage.2009.10.003

[46] Ryali, S., Supekar, K., Abrams, D.A., Menon, V.: Sparse logistic regression for whole-brain classification of fMRI data. NeuroImage 51(2), 752–764 (2010) 10.1016/j.neuroimage.2010.02.040

[47] Eshraghian, J.K., Ward, M., Neftci, E., Wang, X., Lenz, G., Dwivedi, G., Bennamoun, M., Jeong, D.S., Lu, W.D.: Training spiking neural networks using lessons from deep learning. Proceedings of the IEEE 111(9), 1016–1054 (2023)

[48] Paşca, S.P.: The rise of three-dimensional human brain cultures. Nature 553(7689), 437–445 (2018)

[49] Tanveer, M.S., Patel, D., Schweiger, H.E., Abu-Bonsrah, K.D., Watmuff, B., Azadi, A., Pryshchep, S., Narayanan, K., Puleo, C., Natarajan, K., Mostajo-Radji, M.A., Kagan, B.J., Wang, G.: Starting a synthetic biological intelligence lab from scratch. Patterns 6(5) (2025) 10.1016/j.patter.2025.101232. Publisher: Elsevier

[50] Kagan, B.J., Habibollahi, F., Watmuff, B., Azadi, A., Doensen, F., Loeffler, A., Byun, S.H., Servais, B., Desouza, C., Abu-Bonsrah, K.D., Rosbo, N.: Harnessing Intelligence from Brain Cells In Vitro. The Neuroscientist 31(5), 536–555 (2025) 10.1177/10738584251321438. Publisher: SAGE Publications Inc STM

[51] Habibollahi, F., Kagan, B.J., Burkitt, A.N., French, C.: Critical dynamics arise during structured information presentation within embodied in vitro neuronal networks. Nature Communications 14(1), 5287 (2023) 10.1038/s41467-023-41020-3

[52] Tenderra, R.M., Theves, S.: Human intelligence relates to neural measures of cognitive map formation. Cell Reports 44(8), 116033 (2025) 10.1016/j.celrep.2025.116033

[53] Aaser, P., Knudsen, M., Ramstad, O.H., Wijdeven, R., Nichele, S., Sandvig, I., Tufte, G., Stefan Bauer, U., Halaas, , Hendseth, S., Sandvig, A., Valderhaug, V.: Towards making a cyborg: A closed-loop reservoir-neuro system. In: Proceedings of the 14th European Conference on Artificial Life ECAL 2017, pp. 430–437. MIT Press, Lyon, France (2017). 10.7551/ecala072. https://www.mitpressjournals.org/doi/abs/10.1162/isala072

[54] Lindell, T.A.E., Ramstad, O.H., Sandvig, I., Sandvig, A., Nichele, S.: Chaotic Time Series Prediction in Biological Neural Network Reservoirs on Microelectrode Arrays. In: 2024 International Joint Conference on Neural Networks (IJCNN), pp. 1–10. IEEE, Yokohama, Japan (2024). 10.1109/IJCNN60899.2024.10650567. https://ieeexplore.ieee.org/document/10650567/

[55] Sono, Y., Yamamoto, H., Nishi, Y., Sumi, T., Sato, Y., Hirano-Iwata, A., Katori, Y., Sato, S.: Online supervised learning of temporal patterns in biological neural networks under feedback control. Proceedings of the National Academy of Sciences 123(11), 2521560123 (2026) 10.1073/pnas.2521560123

[56] Valverde, S.: Breakdown of Modularity in Complex Networks. Frontiers in Physiology 8 (2017) 10.3389/fphys.2017.00497. Publisher: Frontiers

[57] Wagenaar, D.A., Madhavan, R., Pine, J., Potter, S.M.: Controlling bursting in cortical cultures with closed-loop multi-electrode stimulation 25(3), 680–688 10.1523/JNEUROSCI.4209-04.2005

[58] Maass, W., Natschläger, T., Markram, H.: Real-Time Computing Without Stable States: A New Framework for Neural Computation Based on Perturbations. Neural Computation 14(11), 2531–2560 (2002) 10.1162/089976602760407955

[59] Bullmore, E., Sporns, O.: The economy of brain network organization. Nature Reviews Neuroscience 13(5), 336–349 (2012) 10.1038/nrn3214. Publisher: Nature Publishing Group

[60] Buonomano, D.V., Merzenich, M.M.: Temporal Information Transformed into a Spatial Code by a Neural Network with Realistic Properties. Science 267(5200), 1028–1030 (1995) 10.1126/science.7863330

[61] Ciampi, L., Iannello, L., Tonelli, F., Lagani, G., Di Garbo, A., Cremisi, F., Amato, G.: Neuro-Inspired Visual Pattern Recognition via Biological Reservoir Computing. arXiv. Version Number: 1 (2026). 10.48550/ARXIV.2602.05737. https://arxiv.org/abs/2602.05737

[62] Gorban, A.N., Tyukin, I.Y.: Blessing of dimensionality: mathematical foundations of the statistical physics of data. Philosophical Transactions of the Royal Society A: Mathematical, Physical and Engineering Sciences 376(2118), 20170237 (2018) 10.1098/rsta.2017.0237

[63] Hastie, T., Tibshirani, R., Friedman, J.: The Elements of Statistical Learning: Data Mining, Inference, and Prediction, 2nd edn. Springer Series in Statistics. Springer, New York, NY (2009). 10.1007/978-0-387-84858-7

[64] Benjamini, Y., Hochberg, Y.: Controlling the false discovery rate: A practical and powerful approach to multiple testing. Journal of the Royal Statistical Society: Series B (Methodological) 57(1), 289–300 (1995) 10.1111/j.2517-6161.1995.tb02031.x

[65] Cerqueira, V., Torgo, L., Mozetič, I.: Evaluating time series forecasting models: an empirical study on performance estimation methods. Machine Learning 109, 1997–2028 (2020) 10.1007/s10994-020-05910-7

[66] Sun, Y., Li, X., Dalal, K., Xu, J., Vikram, A., Zhang, G., Dubois, Y., Chen, X., Wang, X., Koyejo, S., Hashimoto, T., Guestrin, C.: Learning to (Learn at Test Time): RNNs with Expressive Hidden States (2024). 10.48550/ARXIV.2407.04620

[67] Khajehnejad, M., Habibollahi, F., Khajehnejad, A., Kagan, B.J., Razi, A.: TAVRNN: Temporal Attention-enhanced Variational Graph RNN Captures Neural Dynamics and Behavior. arXiv. arXiv:2410.00665 [q-bio] (2024). http://arxiv.org/abs/2410.00665

[68] Khajehnejad, M., Habibollahi, F., Khajehnejad, A., French, C., Kagan, B.J., Razi, A.: Graph-based representation learning of neuronal dynamics and behavior. Communications Biology (2026) 10.1038/s42003-026-10721-y

[69] Karimi, M., Bahrami, S., Mirshekari, H., Basri, S.M.M., Nik, A.B., Aref, A.R., Akbari, M., Hamblin, M.R.: Microfluidic systems for stem cell-based neural tissue engineering. Lab on a Chip 16(14), 2551–2571 (2016) 10.1039/C6LC00489J

[70] Zhang, Y., Pak, C., Han, Y., Ahlenius, H., Zhang, Z., Chanda, S., Marro, S., Patzke, C., Acuna, C., Covy, J., Xu, W., Yang, N., Danko, T., Chen, L., Wernig, M., Südhof, T.: Rapid Single-Step Induction of Functional Neurons from Human Pluripotent Stem Cells. Neuron 78(5), 785–798 (2013) 10.1016/j.neuron.2013.05.029

[71] Gantner, C.W., Hunt, C.P.J., Niclis, J.C., Penna, V., McDougall, S.J., Thompson, L.H., Parish, C.L.: FGF-MAPK signaling regulates human deep-layer corticogenesis. Stem Cell Reports 16(5), 1262–1275 (2021) 10.1016/j.stemcr.2021.03.014

[72] Rosebrock, D., Arora, S., Mutukula, N., Volkman, R., Gralinska, E., Balaskas, A., Aragonès Hernández, A., Buschow, R., Brändl, B., Müller, F.-J., Arndt, P.F., Vingron, M., Elkabetz, Y.: Enhanced cortical neural stem cell identity through short SMAD and WNT inhibition in human cerebral organoids facilitates emergence of outer radial glial cells 24(6), 981–995 10.1038/s41556-022-00929-5

[73] Takemuro, T., Yamamoto, H., Sato, S., Hirano-Iwata, A.: Polydimethylsiloxane microfluidic films for in vitro engineering of small-scale neuronal networks. Japanese Journal of Applied Physics 59(11), 117001 (2020). Publisher: IOP Publishing

[74] Murota, H., Yamamoto, H., Monma, N., Sato, S., Hirano-Iwata, A.: Precision Microfluidic Control of Neuronal Ensembles in Cultured Cortical Networks. Advanced Materials Technologies 10(4), 2400894 (2025) 10.1002/admt.202400894

[75] Kamata, H., Mukuta, Y., Harada, T.: Fully Spiking Variational Autoencoder. Proceedings of the AAAI Conference on Artificial Intelligence 36(6), 7059–7067 (2022) 10.1609/aaai.v36i6.20665

[76] Jolliffe, I.T.: Principal Component Analysis, 2nd edn. Springer, New York, NY (2002)

[77] Shlens, J.: A tutorial on principal component analysis. arXiv preprint arXiv:1404.1100 (2014)

[78] Cunningham, J.P., Yu, B.M.: Dimensionality reduction for large-scale neural recordings. Nature neuroscience 17(11), 1500–1509 (2014)

[79] Gallego, J.A., Perich, M.G., Miller, L.E., Solla, S.A.: Neural manifolds for the control of movement. Neuron 94(5), 978–984 (2017)

[80] Beggs, J.M., Plenz, D.: Neuronal Avalanches in Neocortical Circuits. The Journal of Neuroscience 23(35), 11167–11177 (2003) 10.1523/JNEUROSCI.23-35-11167.2003

[81] Cocchi, L., Gollo, L.L., Zalesky, A., Breakspear, M.: Criticality in the brain: A synthesis of neurobiology, models and cognition. Progress in Neurobiology 158, 132–152 (2017) 10.1016/j.pneurobio.2017.07.002

