## Supplementary Materials for "Handwritten Digit classification with neural cultures is influenced by neural architecture, network dynamics, and decoding methods"

Table S1: Experimental Classification Accuracies by Cell and Experiment Type: Frequency-Domain (Stimulus Blanked)

| **Cell Type** | **Experiment Type** | **Num Recordings** | **Acc. (Mean ± SD)** | **Acc. (Median)** |
|---|---|---|---|---|
| Organoid Cortical | MNIST | 3 | $33.7 \pm 2.5\%$ | 34.8% |
| Organoid Hippo | MNIST | 5 | $37.9 \pm 3.9\%$ | 38.1% |
| Monolayer Cortical | MNIST | 2 | $40.5 \pm 0.7\%$ | 40.5% |
| Monolayer Hippo | MNIST | 8 | $38.4 \pm 4.3\%$ | 38.2% |
| 60-module Cortical | MNIST | 22 | $46.9 \pm 9.5\%$ | 42.7% |
| 60-module Hippo | MNIST | 5 | $33.9 \pm 9.3\%$ | 38.2% |
| SDK Random Spikes (Control) | MNIST | 3 | $9.7 \pm 0.8\%$ | 9.3% |
| Saline (Control) | MNIST | 3 | $11.5 \pm 1.2\%$ | 11.3% |

Table S2: Experimental Classification Accuracies by Cell and Experiment Type: Frequency-Domain (Stimulus Included)

| **Cell Type** | **Experiment Type** | **Num Recordings** | **Acc. (Mean ± SD)** | **Acc. (Median)** |
|---|---|---|---|---|
| Organoid Cortical | MNIST | 3 | $61.7 \pm 3.8\%$ | 61.6% |
| Organoid Hippo | MNIST | 5 | $63.5 \pm 3.3\%$ | 63.7% |
| Monolayer Cortical | MNIST | 2 | $59.5 \pm 0.8\%$ | 59.5% |
| Monolayer Hippo | MNIST | 8 | $62.2 \pm 1.5\%$ | 62.2% |
| 60-module Cortical | MNIST | 22 | $67.3 \pm 10.0\%$ | 64.5% |
| 60-module Hippo | MNIST | 5 | $58.2 \pm 9.9\%$ | 63.2% |
| SDK Random Spikes (Control) | MNIST | 3 | $9.3 \pm 1.1\%$ | 8.8% |
| Saline (Control) | MNIST | 3 | $44.7 \pm 3.7\%$ | 44.4% |

Table S3: Experimental Classification Accuracies by Cell and Experiment Type: Time Bin (Trial Holdout)

| **Cell Type** | **Experiment Type** | **Num Recordings** | **Acc. (Mean ± SD)** | **Acc. (Median)** |
|---|---|---|---|---|
| Organoid Cortical | MNIST | 3 | $10.7 \pm 6.1\%$ | 12.0% |
| Organoid Hippo | MNIST | 5 | $12.8 \pm 4.1\%$ | 12.0% |
| Monolayer Cortical | MNIST | 2 | $12.0 \pm 0.0\%$ | 12.0% |
| Monolayer Hippo | MNIST | 8 | $9.2 \pm 5.2\%$ | 6.0% |
| 60-module Cortical | MNIST | 22 | $19.2 \pm 8.2\%$ | 19.5% |
| 60-module Hippo | MNIST | 5 | $22.8 \pm 9.7\%$ | 26.0% |
| SDK Random Spikes (Control) | MNIST | 3 | $10.1 \pm 0.2\%$ | 10.1% |
| Saline (Control) | MNIST | 3 | $10.3 \pm 0.8\%$ | 10.3% |

Table S4: Experimental Classification Accuracies by Cell and Experiment Type:
Time Bin (Time Holdout)

| Cell Type | Experiment Type | Num Recordings | Acc. (Mean ± SD) | Acc. (Median) |
|---|---|---|---|---|
| Organoid Cortical | MNIST | 3 | $14.0 \pm 0.0\%$ | 14.0% |
| Organoid Hippo | MNIST | 5 | $28.4 \pm 13.3\%$ | 28.0% |
| Monolayer Cortical | MNIST | 2 | $24.0 \pm 8.5\%$ | 24.0% |
| Monolayer Hippo | MNIST | 8 | $37.5 \pm 27.6\%$ | 31.0% |
| 60-module Cortical | MNIST | 22 | $51.0 \pm 31.4\%$ | 40.0% |
| 60-module Hippo | MNIST | 5 | $74.4 \pm 9.5\%$ | 74.0% |
| SDK Random Spikes (Control) | MNIST | 3 | $5.3 \pm 1.5\%$ | 5.0% |
| Saline (Control) | MNIST | 3 | $8.8 \pm 1.6\%$ | 9.3% |

Table S5: Experimental Classification Accuracies by Cell and Experiment Type:
Frequency-Domain (Single-Trial / One-Shot)

| Cell Type | Experiment Type | Num Recordings | Acc. (Mean ± SD) | Acc. (Median) |
|---|---|---|---|---|
| Organoid Cortical | MNIST | 3 | $27.0 \pm 2.1\%$ | 27.2% |
| Organoid Hippo | MNIST | 5 | $28.7 \pm 2.2\%$ | 28.4% |
| Monolayer Cortical | MNIST | 2 | $30.3 \pm 0.5\%$ | 30.3% |
| Monolayer Hippo | MNIST | 8 | $29.7 \pm 1.3\%$ | 29.8% |
| 60-module Cortical | MNIST | 22 | $30.1 \pm 4.0\%$ | 31.5% |
| 60-module Hippo | MNIST | 5 | $27.7 \pm 7.3\%$ | 29.5% |
| SDK Random Spikes (Control) | MNIST | 3 | $17.2 \pm 13.9\%$ | 9.3% |
| Saline (Control) | MNIST | 3 | $11.5 \pm 1.1\%$ | 11.3% |

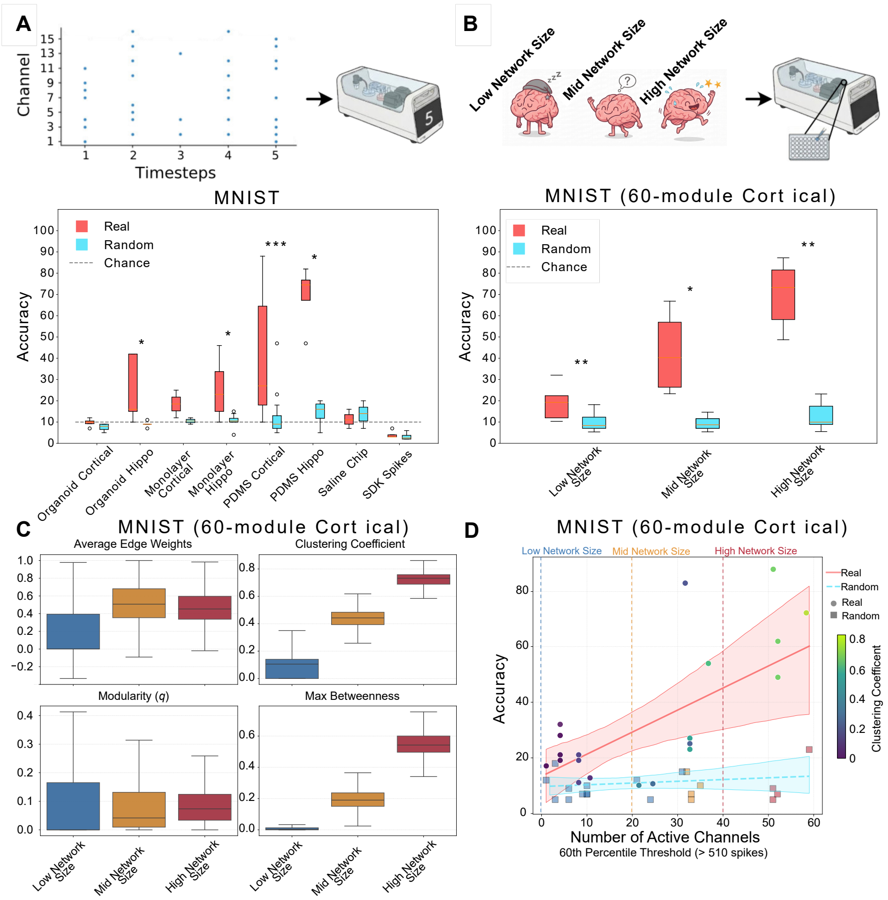


Figure S1: **A**: MNIST Time-Bin results. X-axis: Varying cell types and activity levels. Red reflects results from neuronal cultures, while cyan represents the same data randomly shuffled as a control. Y-axis: accuracy on the MNIST classification task - i.e. how often it was possible to accurately predict which digit was fed into the system just from the system's response. The dashed horizontal line represents chance accuracy. **B**: Accuracy of the 60-module Cortical category from A, separated out as a function of number of active channels. **C**: Functional Connectivity metrics compared with low, medium and high activity from panels A and B. Outliers have been excluded from plotting. **D**: Correlations between real and random accuracy when compared to functional network size (number of active channels), for 60-module Cortical category, coloured by Clustering Coefficient from panel C. Horizontal dotted lines reflect low medium and high network size thresholds. * $p < 0.05$, ** $p < 0.01$, *** $p < 0.001$.

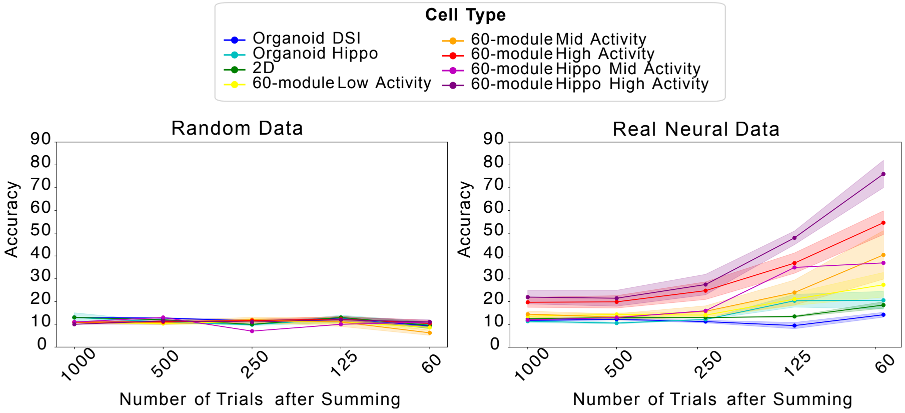


Figure S2: Comparison of accuracy with various summations of individual trials in the Time-Bin (Time Holdout) decoding condition. For n = 1000, spikes from each individual trial were used to calculate accuracy. For n = 500, spikes from every second trial were summed to calculate accuracy, and so on. A higher number of trial summations lead to less variability in cellular activity, and therefore higher overall accuracy.

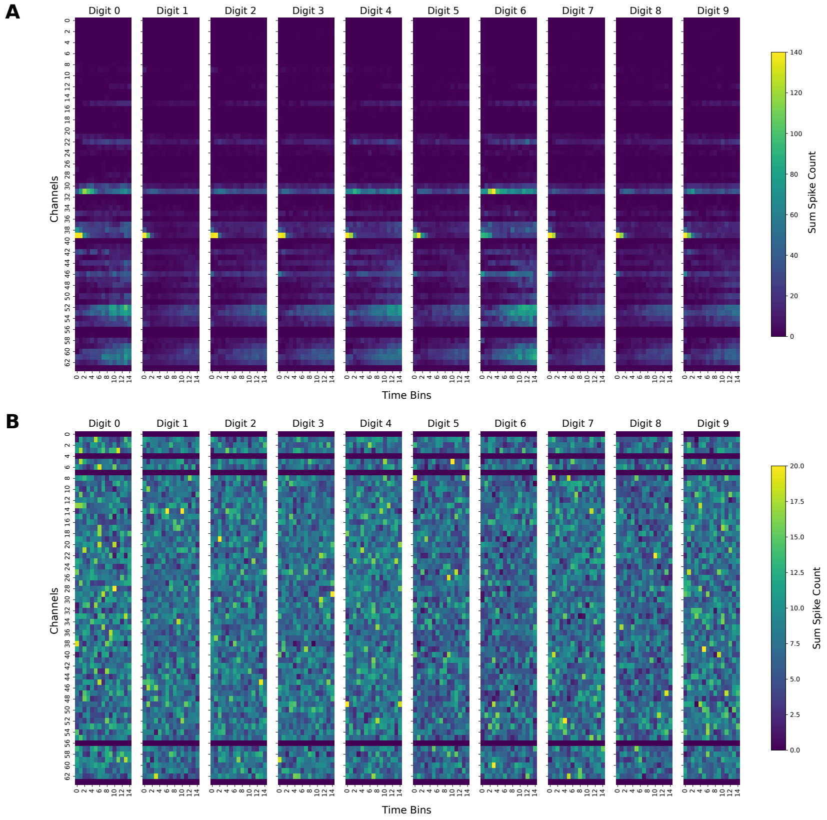


Figure S3: Neural activity as a response to different digit inputs in MNIST. The x-axis the response window of 1 second post stimulation, split into 15 time bins. The y-axis is the Micro-Electrode Array (MEA) channels. **A** Summed spike counts of recorded neural data, per digit. **B** Summed spike counts of randomly shuffled control data, per digit.

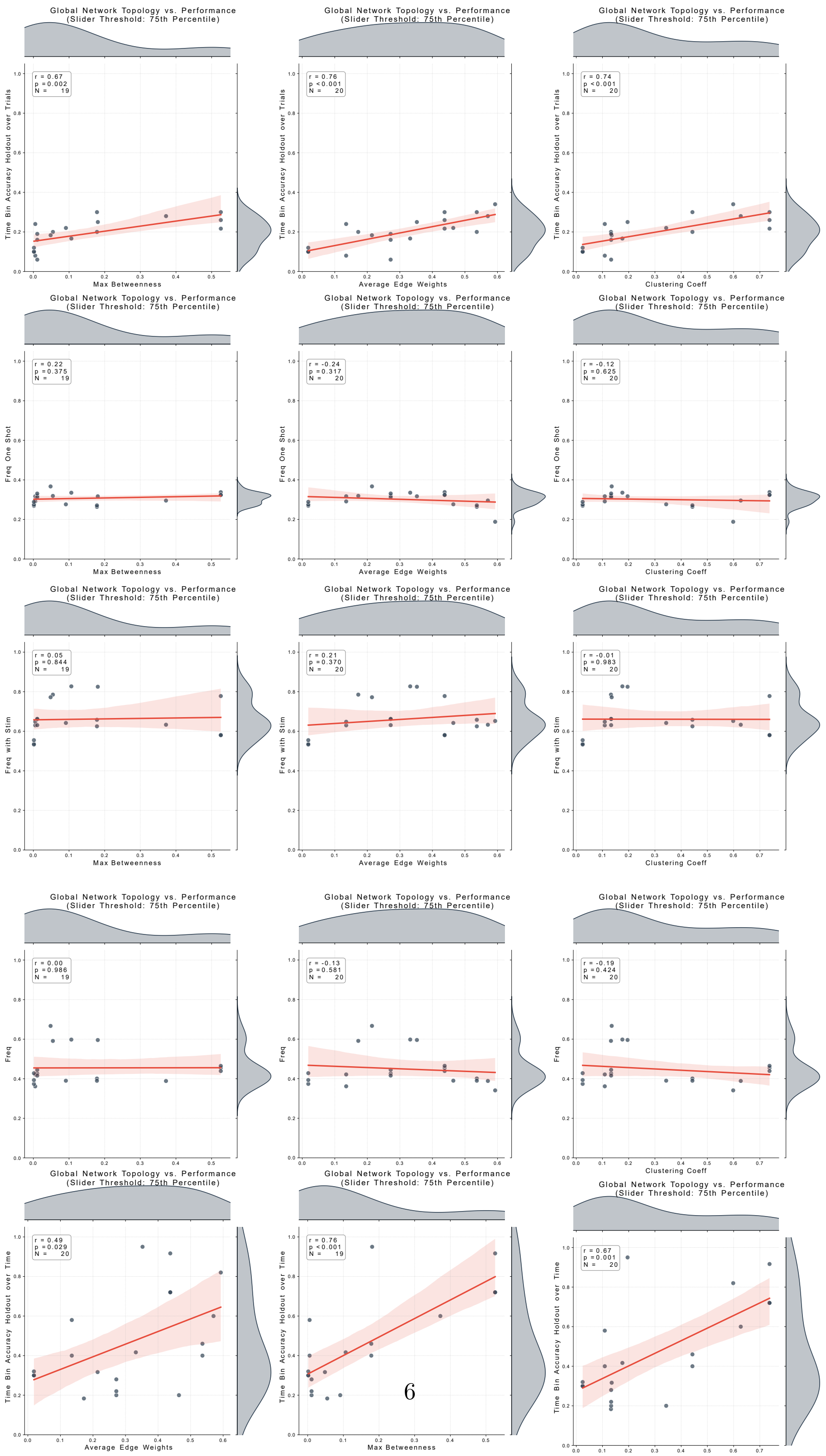




Figure S4: Functional Connectivity metrics vs classification accuracy.

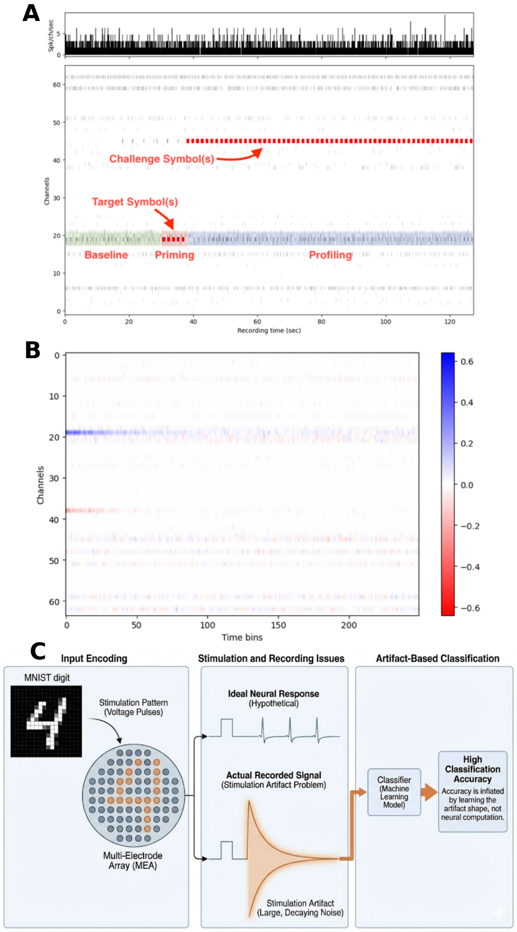


Figure S5: The impact of electrical stimulation artifacts on machine learning classification in MEA recordings. (**A**) Experimental paradigm and recorded activity across 64 channels. The raster plot and top histogram illustrate the recording timeline (~125 seconds), divided into Baseline, Priming, and Profiling phases. Red markers indicate the application of specific “Target” and “Challenge” stimulation symbols. (**B**) A spatiotemporal heatmap (e.g., signal amplitude or classifier weights) across channels and time bins. Distinct horizontal bands align with the stimulation channels (e.g., around channels 19 and 38), indicating a strong spatial footprint linked to the stimulation rather than distributed neural activity. (**C**) Conceptual schematic of the artifact-driven classification problem. When spatial inputs (e.g., MNIST digits) are encoded as voltage pulses on an MEA, the actual recorded signal is dominated by large, decaying stimulation artifacts rather than the ideal neural response. Consequently, machine learning classifiers can achieve artificially high accuracy by learning the shape and profile of the electrical artifact instead of true neural computation.

**Random accuracy parameter sweep vs baseline random accuracy** To determine the effect of number of bins, bin-size and window size on spurious separability, represented by high random accuracy, we conducted a parameter sweep across these variables across different cell types. We then compared the average values across the sweep with the baseline random results (default parameter values). Results are shown in Table S6. Number of bins was the most impactful parameter, with larger number of bins resulting in higher random accuracy, particularly in higher-activity cell types (such as PDMS Hippo; see also S6).

| | Cell Type | N Runs | Mean Sweep Random | Mean Baseline Random | Mean Δ Random | p-value (Sweep Rand > Base Rand) | Above Chance? |
|---|---|---|---|---|---|---|---|
| 0 | Monolayer | 24 | 0.121125 | 0.084706 | 0.036419 | $4.406156 \times 10^{-17}$ | Yes ($p < 0.05$) |
| 1 | Monolayer (Hippo) | 96 | 0.112719 | 0.095294 | 0.017425 | $1.085206 \times 10^{-27}$ | Yes ($p < 0.05$) |
| 2 | Organoid | 36 | 0.124833 | 0.098039 | 0.026794 | $1.073791 \times 10^{-28}$ | Yes ($p < 0.05$) |
| 3 | Organoid (Hippo) | 60 | 0.113567 | 0.096235 | 0.017331 | $2.318902 \times 10^{-16}$ | Yes ($p < 0.05$) |
| 4 | PDMS | 240 | 0.122462 | 0.101926 | 0.020536 | $1.281054 \times 10^{-16}$ | Yes ($p < 0.05$) |
| 5 | PDMS (Hippo) | 60 | 0.295467 | 0.119224 | 0.176243 | $1.172665 \times 10^{-04}$ | Yes ($p < 0.05$) |

Table S6: Random data significance comparison.

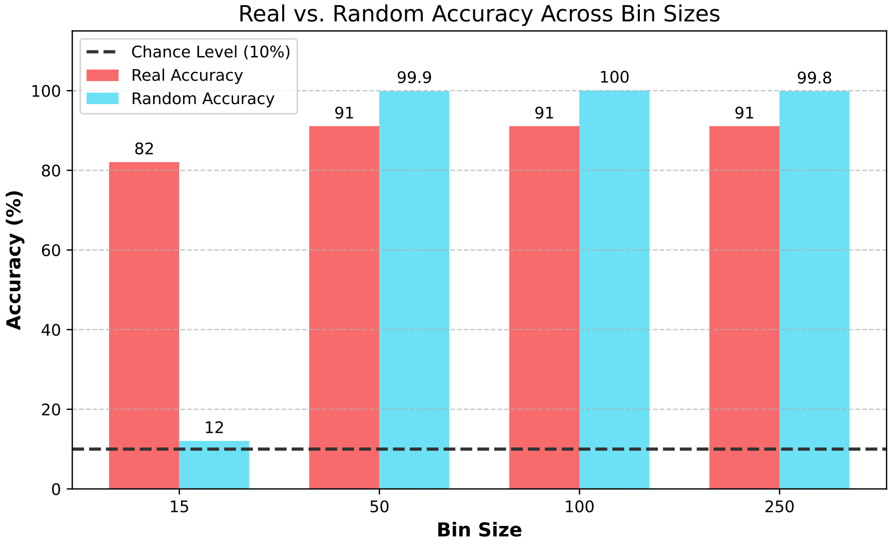


Figure S6: Differences in accuracy and randomised accuracy with increasing time bins. Higher bin sizes result in higher real accuracies, but the trade-off is no-longer relevant controls (close to 100% random accuracy).
